# Pivotal Roles for Ribonucleases in *Streptococcus pneumoniae* Pathogenesis

**DOI:** 10.1101/2021.05.04.442624

**Authors:** Dhriti Sinha, Jacob Frick, Kristen Clemmons, Malcolm E. Winkler, Nicholas R. De Lay

**Affiliations:** Department of Microbiology and Molecular Genetics, McGovern Medical School, University of Texas Health Science Center, Houston, TX 77030, USA; Department of Biology, Indiana University Bloomington, Biology Building; 1001 East Third Street; Bloomington, Indiana 47405, USA; MD Anderson Cancer Center UTHealth Graduate School of Biomedical Sciences, University of Texas Health Science Center, Houston, TX 77030, USA

**Keywords:** RNase Y, polynucleotide phosphorylase, small RNAs, post-transcriptional regulation

## Abstract

RNases perform indispensable functions in regulating gene expression in many bacterial pathogens by processing and/or degrading RNAs. Despite the pivotal role of RNases in regulating bacterial virulence factors, the functions of RNases have not yet been studied in the major human respiratory pathogen *Streptococcus pneumoniae* (pneumococcus). Here, we sought to determine the impact of two conserved RNases, the endoribonuclease RNase Y and exoribonuclease polynucleotide phosphorylase (PNPase), on the physiology and virulence of *S. pneumoniae* serotype 2 strain D39. We report that RNase Y and PNPase are essential for pneumococcal pathogenesis as both deletion mutants showed strong attenuation of virulence in murine models of invasive pneumonia. Genome-wide transcriptomic analysis revealed that nearly 200 mRNA transcripts were significantly up-regulated, whereas the abundance of several pneumococcal sRNAs, including the Ccn (<u>C</u>iaR <u>C</u>ontrolled <u>N</u>oncoding RNA) sRNAs, were altered in the Δ*rny* mutant relative to the wild-type strain. Additionally, lack of RNase Y resulted in pleiotropic phenotypes that included defects in pneumococcal cell morphology and growth *in vitro*. In contrast, Δ*pnp* mutants showed no growth defect *in vitro*, but differentially expressed a total of 40 transcripts including the tryptophan biosynthesis operon genes and numerous 5’-cis-acting regulatory RNAs, a majority of which were previously shown to impact pneumococcal disease progression in mice using the serotype 4 strain TIGR4. Altogether our data suggest that RNase Y exerts a global impact on pneumococcal physiology, while PNPase-mediates virulence phenotypes, likely through sRNA regulation.

**IMPORTANCE:** *Streptococcus pneumoniae* is a notorious human pathogen that adapts to conditions in distinct host tissues and responds to host cell interactions by adjusting gene expression. Ribonucleases (RNases) are key players that modulate gene expression by mediating the turnover of regulatory and protein-coding transcripts. Here, we characterized two highly conserved RNases, RNase Y and PNPase, and evaluated their impact on the S*. pneumoniae* transcriptome for the first time. We show that PNPase influences the levels of a narrow set of mRNAs, but a large number of regulatory RNAs primarily implicated in virulence control, whereas RNase Y has a more sweeping effect on gene expression, altering levels of transcripts involved in diverse cellular processes including cell division, metabolism, stress response, and virulence. This study further reveals that RNase Y regulates expression of genes governing competence by mediating the turnover of <u>C</u>iaR-<u>c</u>ontrolled-<u>n</u>oncoding (Ccn) sRNAs.

## INTRODUCTION

The Gram-positive bacterium *Streptococcus pneumoniae* (pneumococcus) is a common colonizer of the human nasopharynx, where it can remain as a commensal. However, specific signals including host viral infection and environmental and nutritional stress can stimulate *S. pneumoniae* to disperse into other host tissues (1, 2), including the lungs, blood, and brain, and dissemination of pneumococcus into these tissues leads to pneumonia, sepsis, and meningitis, respectively. Pneumococcal infections result in over 1 million deaths annually worldwide (3). *S. pneumoniae* has been shown in murine infection models to have distinct gene expression profiles depending on whether it resides in the blood, brain, nasopharynx, or lungs (4, 5), indicating that it has to adapt to the different conditions in these tissues to survive. Furthermore, pneumococcus also rapidly reprograms its gene expression pattern upon exposure to host cells, such as macrophages (6) and lung epithelial cells (7, 8). To rapidly adapt to changes in their environment, bacteria not only need to modulate the transcription of particular genes, but they must also turnover existing small regulatory RNAs (sRNAs) or mRNAs that encode proteins detrimental under the new set of conditions. Ribonucleases (RNases) control the steady-state levels and turnover of various classes of RNAs (9, 10). In the model Gram-positive bacterium *Bacillus subtilis*, the primary RNase responsible for initiating RNA decay was shown to be RNase Y (11). Depletion of this RNase from *B. subtilis* impacted expression of ≈20% of all open reading frames in its genome (12) and led to a two-fold increase in the half-life of bulk mRNA (11).

RNase Y is the functional, but evolutionarily distinct equivalent of endoribonuclease RNase E of Gram-negative bacteria. RNase Y consists of an N-terminal transmembrane domain followed by a coil-coiled domain, RNA binding KH domain, catalytic HD domain, and a C-terminal domain (12). Like RNase E, RNase Y associates with the membrane (13) and also serves as the organizing component of the RNA degradosome, the central RNA degrading machine in bacteria (12). RNase Y forms this complex by interacting with the RNA helicase CshA, RNases J1 and J2, the glycolytic enzymes phosphofructokinase and enolase, and the exoribonuclease PNPase (12, 14). The dual function RNases J1/J2 exhibit both endo- and 5’-to-3’ exoribonucleolytic activities and are unique to Gram-positive bacteria (15). In *Bacillus subtilis* and *Streptococcus pyogenes*, the decay intermediates resulting from endonucleolytic cleavage are primarily cleared by PNPase, which functions as the major 3’-to-5’ exoribonuclease (16, 17). PNPase has also been shown to significantly impact global mRNA turnover under cold stress in *B. subtilis* and *Staphylococcus aureus*, similar to what has been demonstrated in *E. coli* (18, 19).

Very recent RNA-seq studies in *S. pyogenes* have uncovered the RNA targetomes of both RNase Y and PNPase and further demonstrated that these two proteins work in concert to regulate 5’-regulatory RNA turnover and the stability of polycistronic mRNAs (20). These results are consistent with the previously implicated role of RNase Y in mediating decay of 5’ cis-acting regulatory RNAs (S-adenosylmethionine, T-box, and riboflavin riboswitches) in *S. aureus* and *B. subtilis* (11, 21, 22). Interestingly, results of a recent study that globally examined protein-RNA associations in *S. pneumoniae* via gradient profiling by sequencing (Grad-seq) indicate that PNPase interacts with several small RNAs *in vivo* (23). However, compared to Gram-negative bacteria the detailed mechanisms by which major Gram-positive RNases, such as RNase Y and PNPase, impact sRNA-dependent regulation and *trans*-acting sRNA levels remain largely unknown; but, there has been some evidence for RNase Y-dependent turnover of sRNAs (e.g., RsaA in *S. aureus* and RoxS in *B. subtilis* (24)). Independent studies have further indicated an indirect role of RNase Y in regulating the abundance of two other *trans*-acting sRNAs VR-RNA and FasX in the important Gram-positive pathogens *Clostridium perfringens* and *S. pyogenes*, respectively (25, 26). The extent to which RNase Y orthologs from different species contribute to growth and RNA decay vary considerably (22). These findings further emphasize that various Gram-positives, including pathogens, may employ different mechanistic strategies to mediate RNA decay and processing.

In spite of the crucial roles of RNases in impacting bacterial stress response by altering gene expression, we do not know about the functions of major pneumococcal RNases. In the present work, we report characterization of two conserved RNases, RNase Y and PNPase, in *S. pneumoniae* serotype 2 strain D39. We demonstrate that RNase Y functions as a broadly pleiotropic regulator, whose absence significantly impacts the pneumococcal mRNA transcriptome, growth, virulence, and stability and function of conserved pneumococcal Ccn sRNAs. In contrast, PNPase impacts the abundance of several important transcripts, including riboswitches that were previously implicated in pneumococcal virulence control. Consistently the absence of PNPase resulted in a strong virulence defect *in vivo*, while displaying no obvious phenotypes *in vitro*. Altogether, our work has uncovered for the first time the crucial roles for two well-conserved RNases in regulating pneumococcal physiology and virulence.

## RESULTS

### RNase Y is required for normal pneumococcal growth and cell morphology

Prior studies have shown that deletion of *rny*, the gene encoding RNase Y, from *B. subtilis* and *C. perfringens* caused a drastic reduction in growth, but the effect of removal of this gene on *S. pyogenes* and *S. aureus* growth was modest (25, 27–29). However, deletion of *pnp* led to a cold-sensitive phenotype in *B. subtilis*, similar to what was observed for *Escherichia coli* (30, 31). Therefore, we assessed the effects of clean deletion in *rny* or *pnp* (Table S1) on pneumococcal growth at both optimal (37°C) and lower temperatures (32°C). We found that at 37°C in brain heart infusion (BHI) broth, the Δ*rny* mutant exhibited a significant reduction in growth rate and yield compared to the wild-type (WT) strain (Figs 1A; Table S2). The average doubling time and growth yield for the Δ*rny* mutant was ≈76 min and 0.13 compared with ≈45 min and 0.83 for the WT strain. The observed growth defect of the Δ*rny* mutant was restored by repairing the mutation to the WT allele at the native locus (Fig. 1A and Table S2). We also observed that the growth deficiency of the Δ*rny* mutant became more pronounced in 15- and 25-day old BHI compared to freshly prepared (≤ 5-day old) BHI, whereas the isogenic parental strain grew similarly in fresh and aged BHI (Figs. S1A and S1B; Table S2). In contrast, the Δ*pnp* mutant grew like the WT strain in BHI broth at 37°C (Fig. 1A, Figs. S1A and S1B). In contrast to *E. coli* and *B. subtilis* (30, 31), the pneumococcal Δ*pnp* mutant did not show a cold-sensitive (CS) phenotype (Fig. 1B) at 32°C, the lowest temperature at which *S. pneumoniae* D39 grows well. Finally, the Δ*rny* mutant was not cold-sensitive, as the relative growth rate differences between the WT and Δ*rny* mutant were not significantly different at 32°C compared to 37°C (Fig. 1B; see Fig. S1C and Table S2).

**Figure 1:**
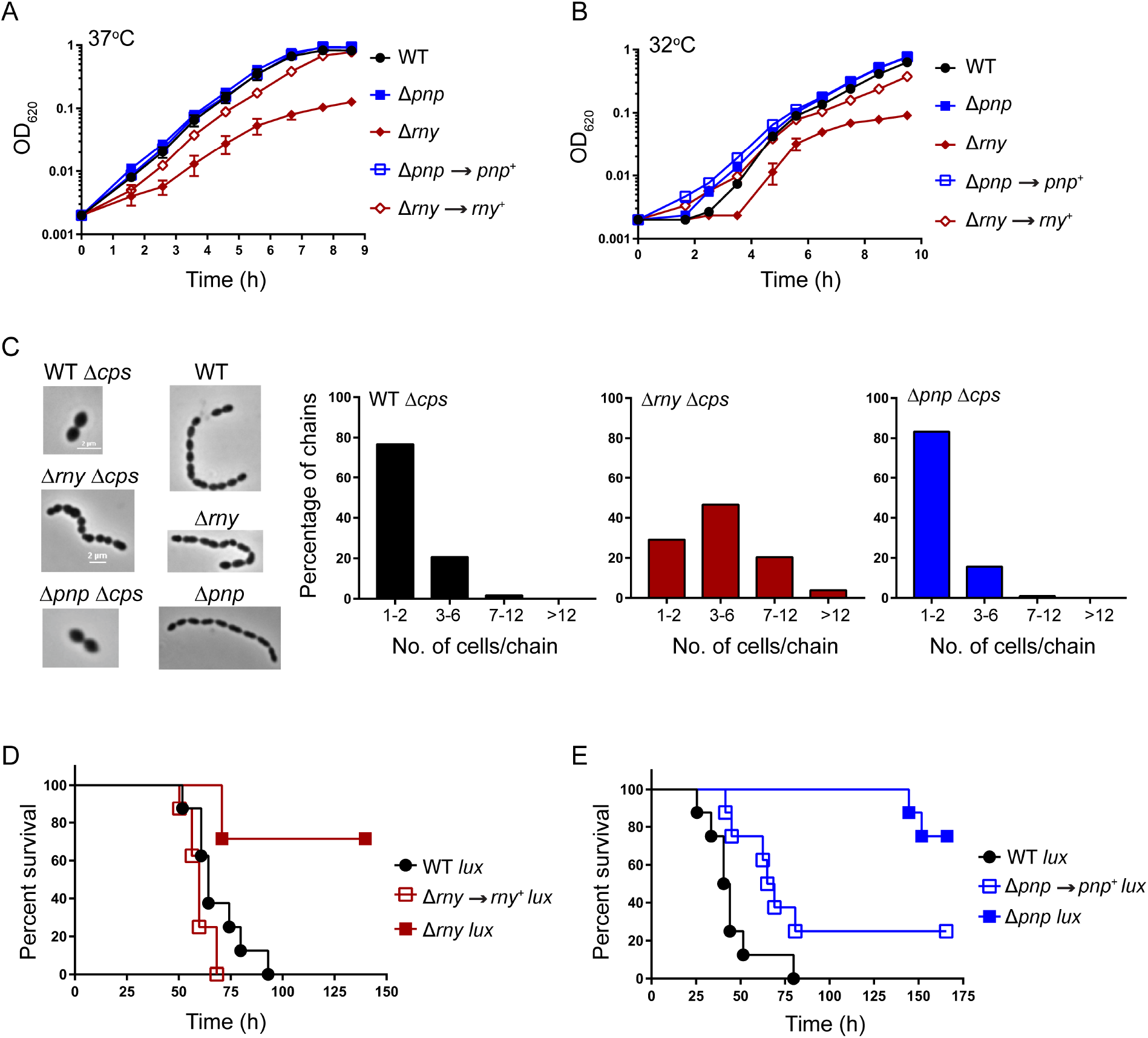
Phenotypes of Δ*rny* and Δ*pnp* mutants. (A, B) Growth characteristics of encapsulated D39 parent strain (IU1781) and isogenic mutant strains, Δ*rny* (NRD10092), Δ*pnp* (IU4883), Δ*rny//rny^+^* (NRD10305), and repaired strain Δ*pnp//pnp^+^* (NRD10303), grown statically at 37°C and at 32°C in 15-day-old BHI broth in an atmosphere of 5% CO_2_. Growth curves represent data from three independent replicates for each strain at 37°C or 32°C. Average growth rates and growth yields are listed in Table S2. (C) Representative phase-contrast images of the D39 wild-type strain (WT; IU1781) and its derived mutant strains Δ*cps* (IU1824), Δ*rny* (NRD10092), Δ*pnp* (IU4883), Δ*cps* Δ*rny* (NRD10109), and Δ*cps* Δ*pnp* (NRD10108), in early exponential growth phase. Distribution of chain lengths were based on 100 to 110 chains from at least two independent cultures of each strain. (D, E) Survival curve analysis showing disease progression in an invasive murine model of pneumonia. ICR male mice were inoculated intranasally with ≈10^7^ CFUs in 50 µL inocula of the D39 parent expressing a *lux* luminescence cassette (D39 Tn*4001 luxABCDE*; IU1918) or isogenic mutants (Δ*rny* Tn*4001 luxABCDE*, IU6838; Δ*pnp* Tn*4001 luxABCDE*, IU6622; *rny*^+^ Tn*4001 luxABCDE*, IU7152; and *pnp*^+^ Tn*4001 luxABCDE*, IU7154). Eight animals were infected per strain and disease progression was followed in real time by survival curve analysis (*Materials and Methods*). Survival curves were analyzed by Kaplan- Meier statistics and log-rank tests to determine P-values.

To gain insight into the growth impairment of Δ*rny* mutants, we examined cells from early exponential phase cultures (OD_620_ ≈0.1-0.15) of the WT and Δ*rny* mutant by phase contrast microscopy. We found that the Δ*rny* mutant exhibits significant morphological defects. Occasionally Δ*rny* mutants formed minicells at ends or in the middle of a chain, indicating a possible cell division defect (Fig. 1C). The abnormalities in cell morphology that we observed in the encapsulated Δ*rny* mutant were even more pronounced in a Δ*cps* mutant lacking capsule (Fig. 1C). This observation is consistent with previous findings that capsule tends to dampen pneumococcal cell shape and division phenotypes (32). In addition, the Δ*rny* Δ*cps* mutant formed longer chains comprising of 4-12 cells/chain compared to the WT parent, which were mainly diplococci (Fig. 1C). Finally, we did not observe any morphological differences between the Δ*pnp* mutant and the WT parent in either *cps*^+^ or Δ*cps* background (Fig. 1C). We conclude that RNase Y, but not PNPase is required for *S. pneumoniae* D39 normal growth and cell morphology.

### RNase Y and PNPase are important pneumococcal virulence factors

Lack of RNase Y in *S. pyogenes* and *S. aureus* resulted in virulence attenuation (28, 33, 34). Therefore, we determined the consequences of the *rny* and *pnp* deletions on *S. pneumoniae* D39 pathogenesis using a murine invasive pneumonia model (see Materials and Methods). Both the Δ*rny* and Δ*pnp* mutants were substantially attenuated for virulence compared to the WT parent (Fig. 1D and 1E). 75% and 87% of the mice inoculated with the Δ*pnp* or Δ*rny* mutant, respectively, survived the entire course of the experiment (≈170 h), whereas the median survival time for the WT parent strain ranged between 42 h to 64 h (Fig. 1D, 1E, and S2). To determine if the attenuated virulence observed in each case was correlated with loss of *rny* or *pnp* function, we repaired Δ*rny* or Δ*pnp* back to the *rny*^+^ or *pnp*^+^ allele, respectively, by allelic exchange (Materials and Methods). The *rny*^+^ and *pnp^+^* repaired strain displayed a median survival time of 60 h and 67 h, respectively, indicative of full virulence. Taken together, we conclude that both RNase Y and PNPase contribute to pneumococcal pathogenesis.

### RNase Y and PNPase impact the pneumococcal mRNA transcriptome differently

To identify target transcripts of RNase Y or PNPase that influence pneumococcal physiology, next, we compared the genome-wide transcriptome profiles of Δ*rny* or Δ*pnp* mutant relative to the WT parent grown in matched batches of BHI broth at 37°C in an atmosphere of 5% CO_2_ using mRNA-sequencing (mRNA-seq) analysis (see Materials and Methods). mRNA-seq of the Δ*rny* mutant revealed that 185 transcripts were significantly up-regulated compared to the WT parent strain, using a cutoff >1.8-fold change and a *P* value adjusted for multiple testing (*P*_adj_) <0.05. In contrast, only 28 genes were significantly down-regulated in the Δ*rny* mutant compared to the WT strain (Table 1 and Fig. 2A). The up-regulated transcripts encode proteins that are involved in diverse cellular functions including translation, transcription, transport and metabolism of carbohydrates, amino acids, nucleotides, coenzymes and inorganic ions, cell wall and envelope biogenesis, and stress response (Table 1). In particular, several transcripts that were up-regulated in the Δ*rny* mutant are under the regulatory control of the WalRK, LiaFSR, PnpRS, or CiaRH two-component systems (TCS) (Table 1). Notably, relative transcript abundance for genes encoding important cell division and cell wall proteins, including *mapZ, cozE, gpsB, lytB, licB, licC, licA, tarI, tarJ, spd_0703, and spd_0104*, were increased by ≈2-5-fold in the Δ*rny* strain. Lack of RNase Y also increased the relative expression of genes involved in stress response (*clpL,* 9-fold; *dnaK,* 6-fold; *dnaJ,* 3-fold; *hptX,* 2-fold) and *pavB* (≈2-fold), which encodes a fibronectin-binding protein involved in pneumococcal virulence.

**Figure 2:**
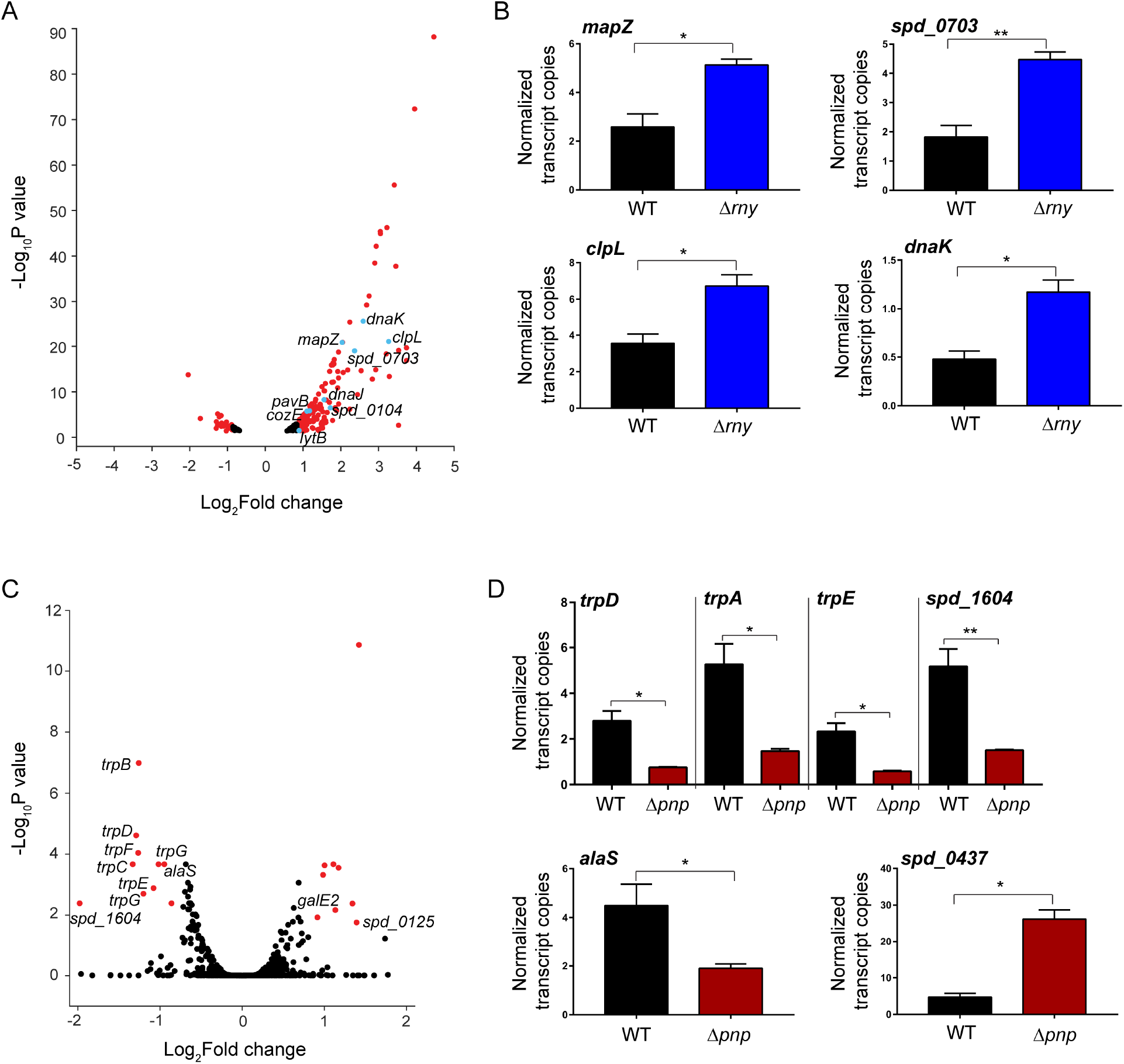
Impact of RNase Y and PNPase on mRNA transcriptome of *S. pneumoniae* D39. (A, C) Volcano plot showing genome-wide changes in mRNA transcript levels in a Δ*rny* mutant (A) or Δ*pnp* mutant (C) relative to the D39 parent strain. RNA was extracted from exponentially growing cultures of the WT D39 parent (IU3116) and isogenic mutants, Δ*rny* mutant (IU5504) or Δ*pnp* mutant (IU5498), in triplicate and analyzed by mRNA-seq as described in *Materials and Methods*. Red and blue dots represent genes with relative transcript changes >1.8-fold as the cut- off (Log_2_fold change =0.85), with an adjusted P-value cut-off <0.05. Relative transcript level changes of genes below the cut-off values are considered to be insignificant and are color coded in black. The X-axis represents gene-fold changes, and the Y-axis represents corresponding P- values plotted on a logarithmic scale. mRNAs that were significantly up-regulated or down- regulated in the Δ*rny* mutant or Δ*pnp* mutant compared to the parent are listed in Table 1 and Table 2, respectively. (B, D) dd-PCR analysis was used to determine copy numbers of indicated transcripts in a wild-type D39 parent (WT; IU1781) and isogenic mutants (Δ*rny*, NRD10092; Δ*pnp*, IU4883). Transcript numbers were normalized to the 16S transcript number, which served as the internal control. Data and error bars represent the means (± SEM) of at least three independent experiments and asterisks (*) and (**) indicate P<0.05 and P<0.01, respectively.

**Table 1:**
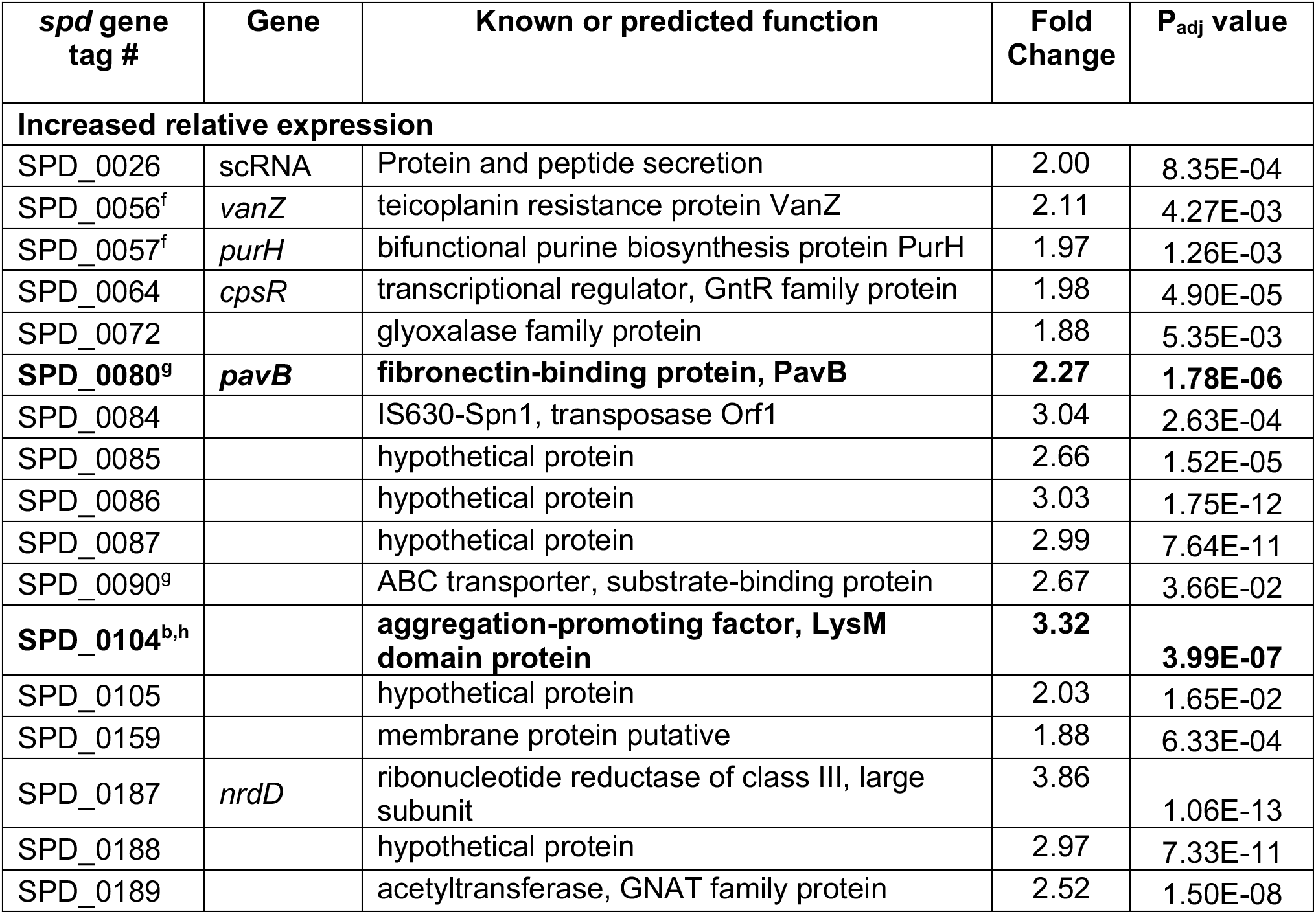
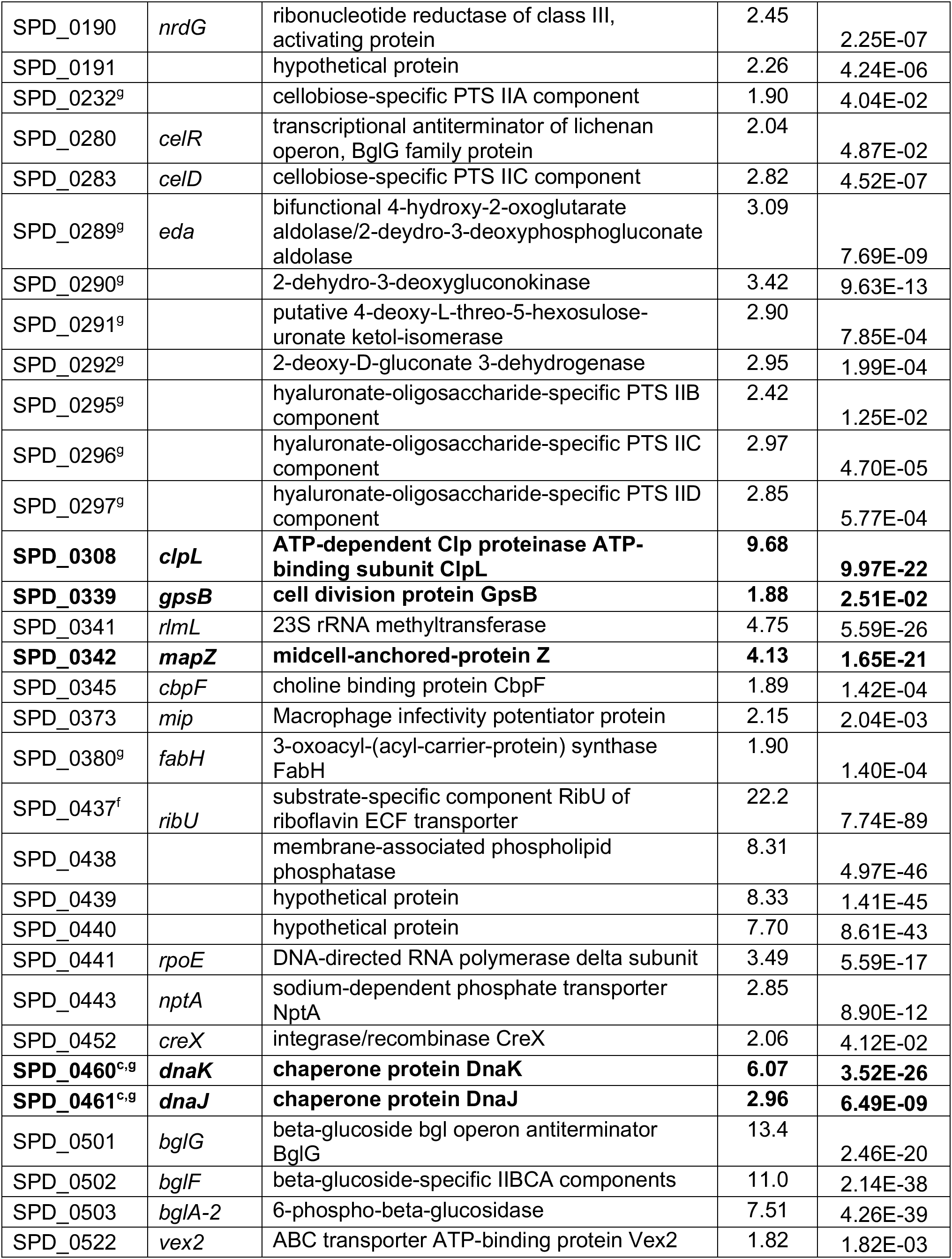
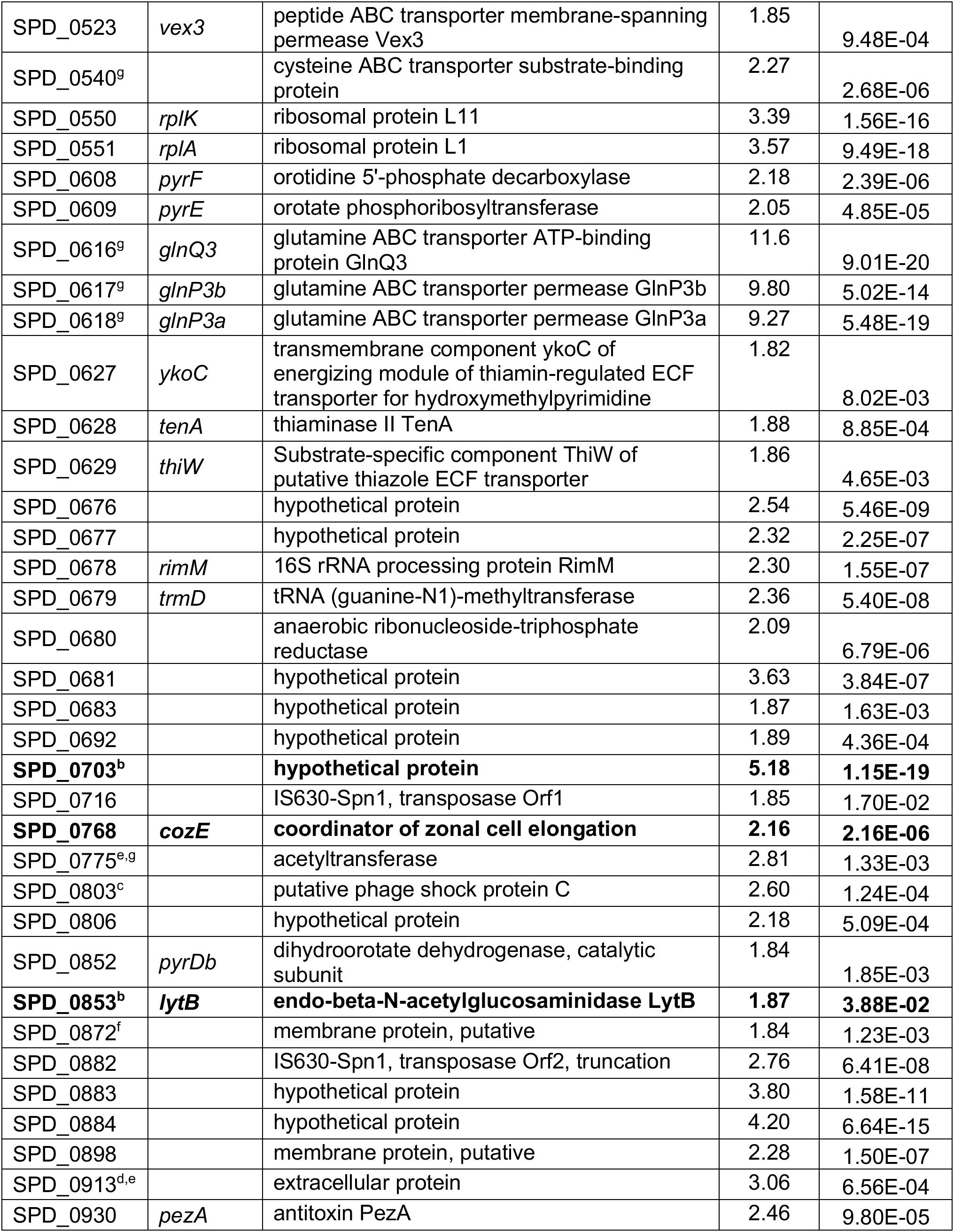
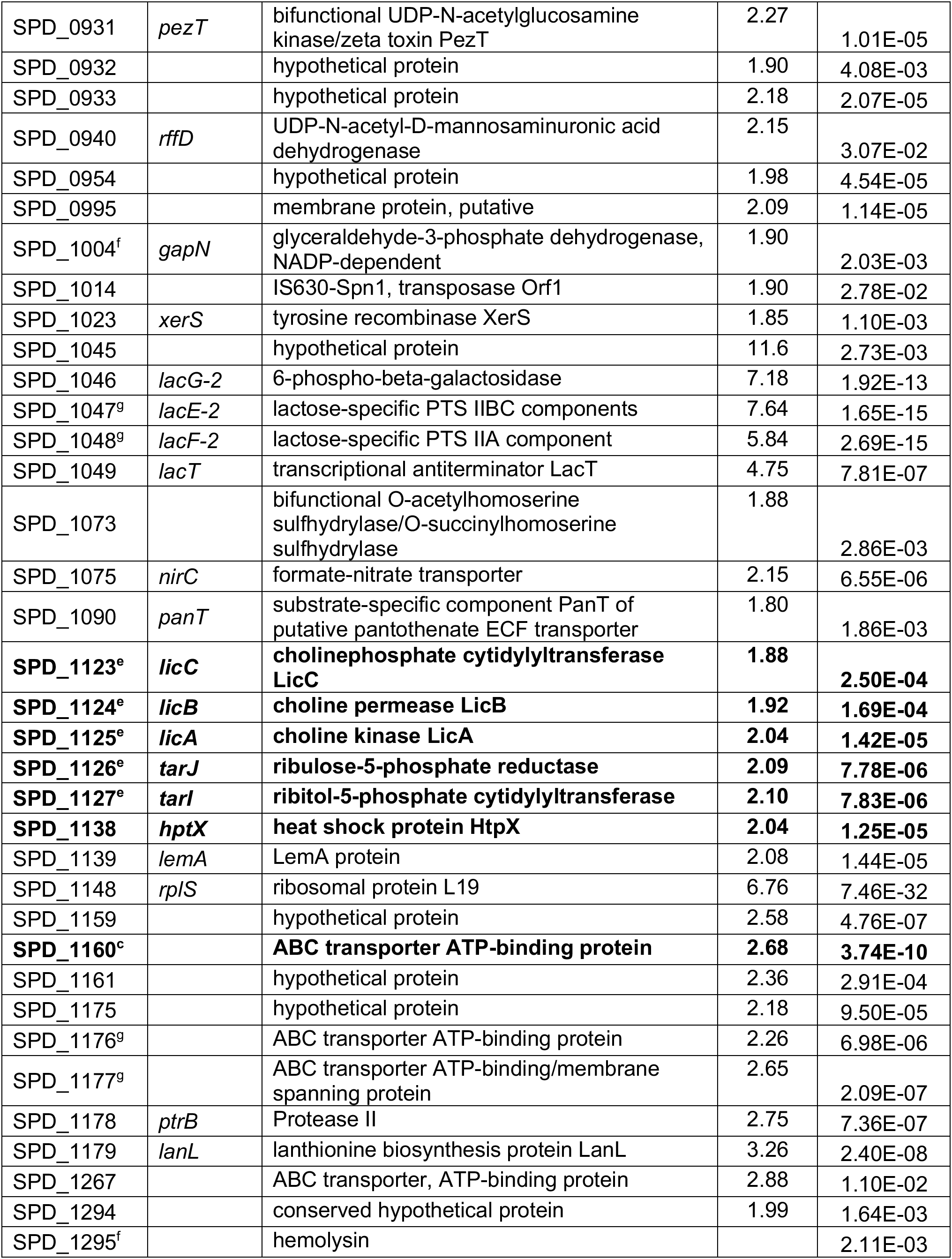
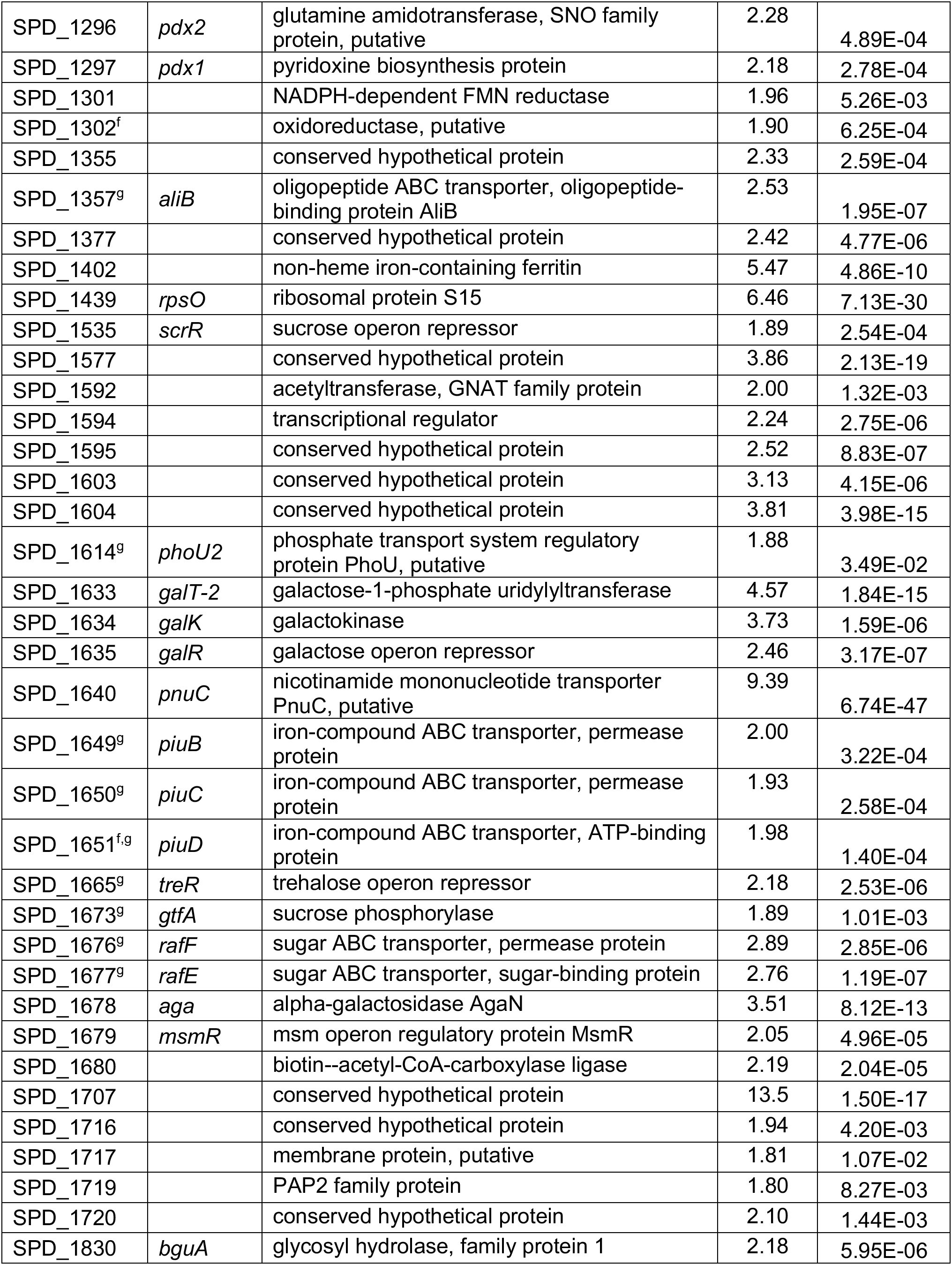
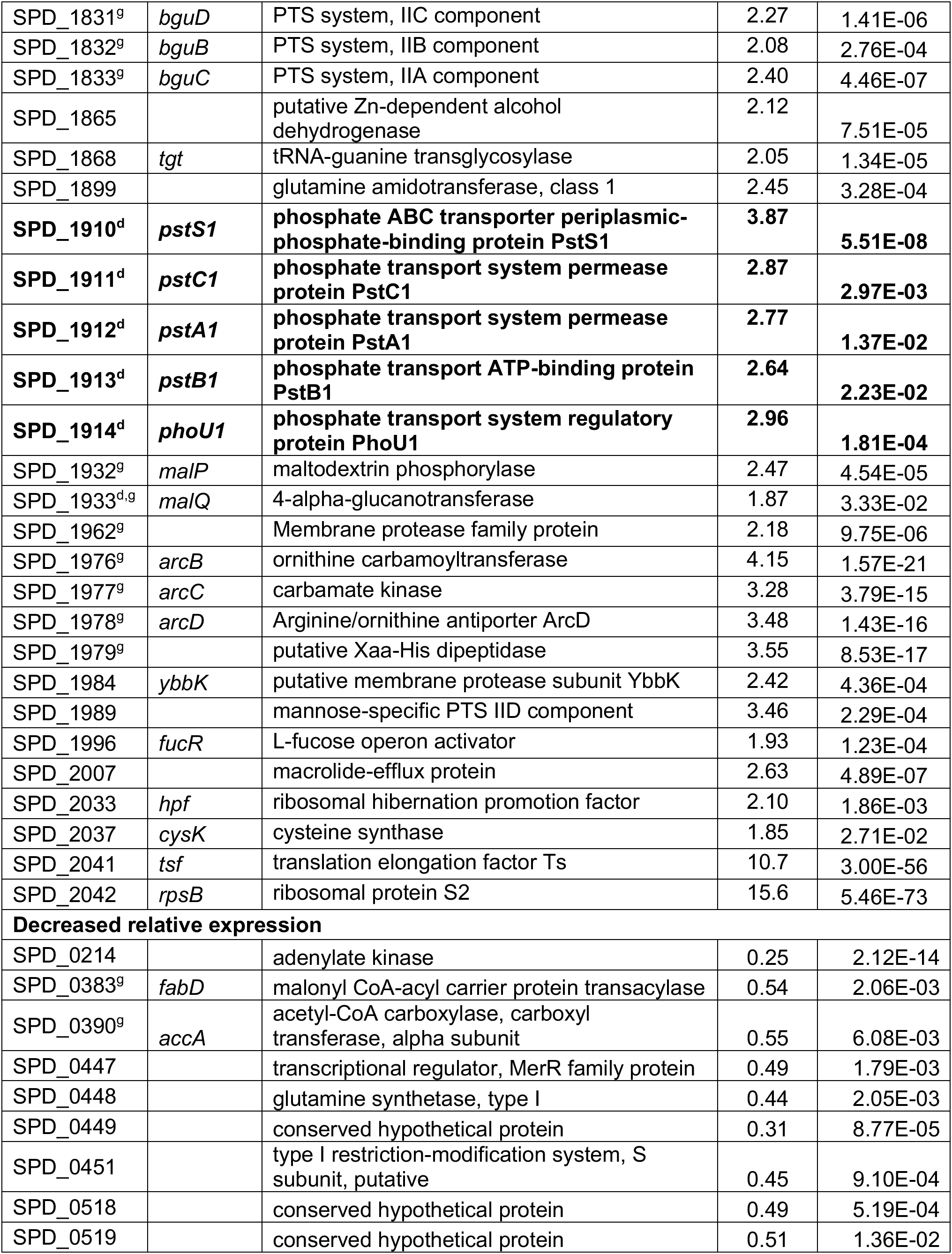
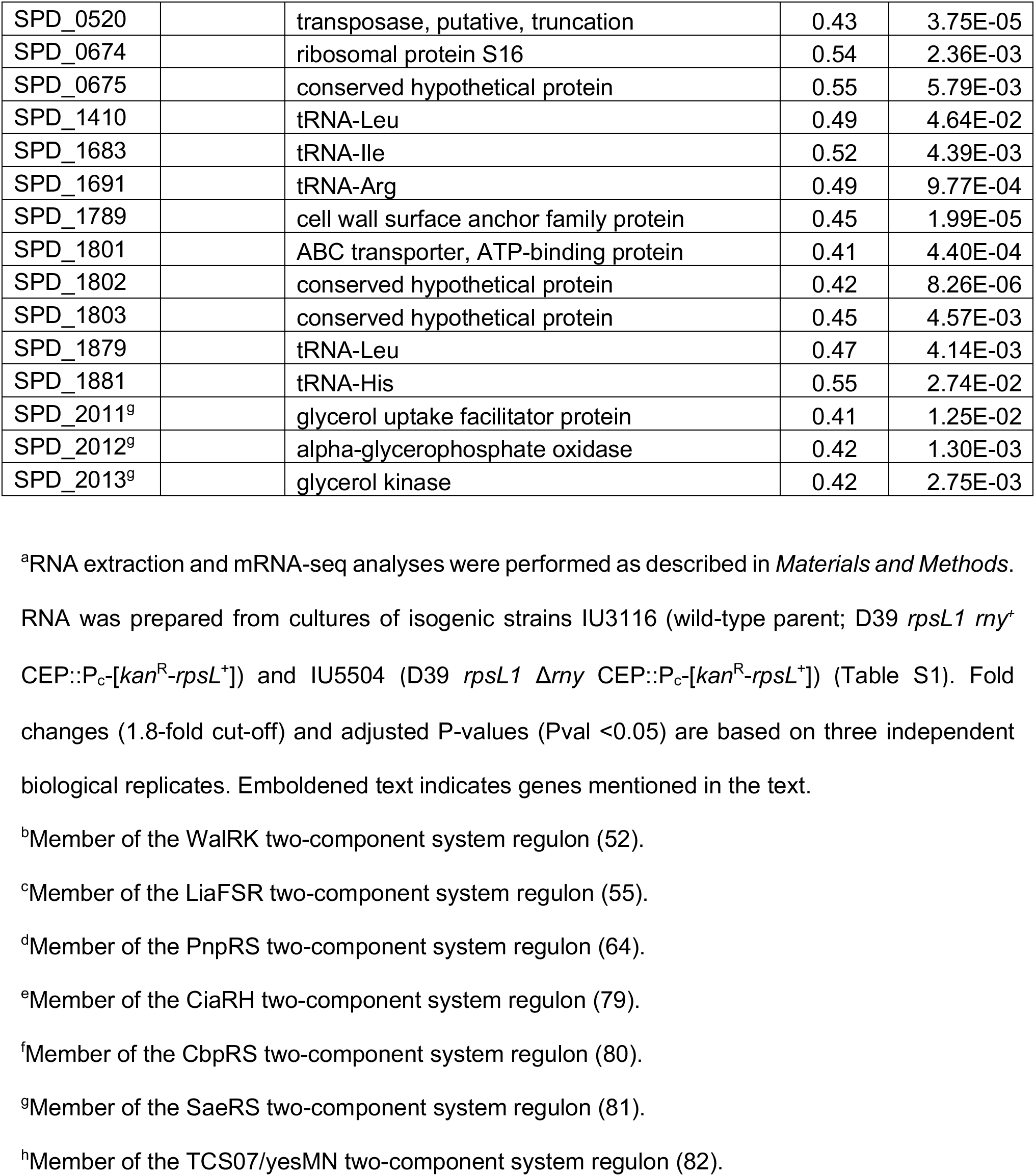
Genes showing changes in relative mRNA transcript amounts in a Δ*rny* mutant compared to the *rny* ^+^ parent strain during exponential growth in BHI broth^a^

**Table 2:** Genes showing changes in relative mRNA transcript amounts in a Δ*pnp* mutant compared to the *pnp* ^+^ parent strain during exponential growth in BHI broth^a^

| *spd* gene tag # | Gene | Known or predicted function | Fold Change | P<sub>adj</sub> value |
| --- | --- | --- | --- | --- |
| **Increased relative expression** |  |  |  |  |
| SPD_0125 |  | Hypothetical protein | 2.62 | 1.72E-02 |
| SPD_0437<sup>b</sup> | *ribU* | Substrate-specific component RibU of riboflavin ECF transporter | 4.16 | 1.33E-36 |
| SPD_0771<sup>b</sup> | *fruR* | Transcriptional repressor of the fructose operon | 2.00 | 2.33E-04 |
| SPD_0975<sup>b</sup> | *radC* | DNA repair protein RadC | 2.00 | 4.75E-04 |
| SPD_1579 |  | Hypothetical protein | 2.53 | 4.13E-03 |
| SPD_1586<sup>b</sup> |  | Multiple sugar metabolism operon regulatory protein | 2.68 | 1.39E-11 |
| SPD_1588 |  | Hypothetical protein | 1.89 | 1.18E-02 |
| SPD_1612<sup>b</sup> | *galE-2* | UDP-glucose 4-epimerase | 2.19 | 6.74E-03 |
| SPD_1707 |  | Hypothetical protein | 2.25 | 2.79E-04 |
| SPD_1792 |  | Hypothetical protein | 2.16 | 2.13E-04 |
| **Decreased relative expression** |  |  |  |  |
| SPD_1216<sup>d</sup> | *alaS* | Alanyl-tRNA synthetase | 0.52 | 2.13E-04 |
| SPD_1596<sup>bc</sup> | *trpA* | Tryptophan synthase alpha chain | 0.49 | 2.13E-04 |
| SPD_1597<sup>c</sup> | *trpB* | Tryptophan synthase beta chain | 0.42 | 1.03E-07 |
| SPD_1598<sup>c</sup> | *trpF* | Phosphoribosylanthranilate isomerase | 0.42 | 9.06E-05 |
| SPD_1599<sup>bc</sup> | *trpC* | Indole-3-glycerol phosphate synthase | 0.40 | 2.13E-04 |
| SPD_1600<sup>bc</sup> | *trpD* | Anthranilate phosphoribosyltransferase | 0.41 | 2.42E-05 |
| SPD_1601<sup>bc</sup> | *trpG* | Bifunctional anthranilate synthase | 0.44 | 1.97E-03 |
| SPD_1602<sup>bc</sup> | *trpE* | Anthranilate synthase, amidase component | 0.47 | 1.29E-03 |
| SPD_1604 |  | Hypothetical protein | 0.25 | 4.06E-03 |
| SPD_2012 | *glpO* | Alpha glycerophosphate oxidase | 0.55 | 4.06E-03 |

<sup>a</sup>RNA extraction and mRNA-seq analyses were performed as described in *Materials and Methods*.
<sup>b</sup>Role in virulence according to Tn-seq studies in TIGR4 (36).
<sup>c</sup>Member of the tryptophan (*trp*) biosynthesis operon.
<sup>d</sup>Likely essential gene according to Tn-seq studies in TIGR4 (36).

Deletion of *pnp* had substantially less impact on relative mRNA transcript amounts, with significant changes in abundance of only twenty transcripts (Table 2 and Fig. 2C). Interestingly, a majority of mRNA transcripts that were differentially regulated in the Δ*pnp* mutant were shown by a previous Tn-seq screen of serotype 4 strain TIGR4 to be important for colonization of the nasophayrnx and/or infection of the lungs in murine infection models (35) (Table 2). In particular, the relative abundance of transcripts corresponding to the tryptophan biosynthesis operon (*trpABCDEFG*), including the upstream gene *spd_1604*, were maximally down-regulated by ≈3- 4-fold. In addition, the relative transcript amount of *alaS*, which encodes alanyl-tRNA-synthetase, was down-regulated by ≈2-fold in the Δ*pnp* mutant compared to the WT strain (Table 2; Fig. 2C). Results from mRNA-seq analyses were confirmed by reverse transcriptase droplet digital PCR (RT-ddPCR) as described in *Material and Methods*. Consistent with the RNA-seq results, the relative transcript amounts of *mapZ* (≈2-fold), *spd_0703* (≈3-fold), *clpL* (≈4-fold), and *dnaK* (≈2-fold) increased in the Δ*rny* mutant compared to WT strain (Fig. 2B). In the Δ*pnp* mutant, RT-ddPCR showed that the relative transcript amounts of *spd_1604*-*trpD*-*trpA*-*trpE* and *alaS* decreased by ≈4-fold and 2.4-fold, respectively, whereas the relative amount of *spd_0437* (*ribU*) increased by ≈6-fold (Fig. 2D), again consistent with the RNA-seq results. Together, these data confirm the relative changes in steady-state mRNA transcript amounts caused by lack of RNase Y or PNPase in *S. pneumoniae*.

### RNase Y and PNPase mediate the sRNA transcriptome of *S. pneumoniae*

Previous studies demonstrate that RNase Y directly and indirectly impacts sRNA levels in several important Gram-positive pathogens (25, 36, 37), whereas, PNPase promotes the stability of some sRNAs and degrade others in *E. coli* (38, 39). A recent Grad-seq study indicates that *S. pneumoniae* PNPase binds to several sRNAs, including CcnA, CcnB, CcnC, CcnD, and Spd_sr34 (23). To further understand how RNases modulate the stability and function of sRNAs expressed by *S. pneumoniae* D39, we sought to identify the sRNAs targeted by RNase Y and PNPase using sRNA-sequencing (sRNA-seq) (see *Materials and Methods*). At least 112 distinct sRNAs have been identified in *S. pneumoniae* D39 (40, 41).

sRNA-seq analysis revealed 11 sRNAs (≈10% of total sRNAs) showed a >1.8-fold change in relative amount between the Δ*rny* mutant and WT strain (Table 3; Fig. 3A). Seven sRNAs were up-regulated in the Δ*rny* mutant compared to the WT strain, whereas only 4 were down-regulated. The putative regulatory RNAs impacted by Δ*rny* fall into all five categories of sRNAs based on their location relative to previously annotated genes in D39 (Fig. 3B). Three of the sRNAs differentially expressed in the Δ*rny* mutant contain regulatory elements; Spd-sr12 and Spd-sr32 contain T-box riboswitches and Spd-sr48 has an L20 leader sequence that regulates the expression of downstream ribosomal genes. Interestingly, among the significantly up-regulated sRNAs in the Δ*rny* mutant are 2 Ccn sRNAs (CcnA and CcnE; Table 3 and Fig. 3A), which are among the five homologous, highly conserved intergenic pneumococcal sRNAs under positive transcriptional control of the CiaR response regulator and function to inhibit competence development via base-pairing with the precursor of the competence stimulatory peptide-encoding mRNA, *comC* (42, 43, 44). Seven out of eleven sRNAs that were differentially expressed in the Δ*rny* mutant relative to the WT strain were experimentally validated using northern blotting. We found that four sRNAs (CcnE, CcnA, Spd-sr12, and Spd-sr32) are significantly upregulated, while for sRNAs Spd-sr100 and Spd-sr116 the annotated full-length transcripts could not be detected in the Δ*rny* mutant (Figs. 4, S3, and S5). However, we did observe that a higher molecular weight band corresponding to Spd-sr116 was increased in abundance only in a Δ*rny* mutant (Fig. S3). Spd-sr108 was the only sRNA for which we observed a significant difference in abundance between the Δ*rny* mutant and WT strain by RNA-seq, but not by northern blot analysis (Table 3, Figs. S3, and S5). In addition to these sRNAs, we probed for 12 additional sRNAs that, were not significantly differentially expressed in the Δ*rny* mutant relative to the WT strain in the RNA-seq analysis. Northern blots revealed that eight of these sRNAs (Spd-sr43, Spd-sr44, Spd-sr73, Spd- sr74, Spd-sr80, Spd-sr83, Spd-sr88, and Spd-sr114) were up-regulated in the Δ*rny* mutant relative to the wild type strain, whereas 4 others (Spd-sr70, Spd-sr54, Spd-sr82, and Spd-sr96) were unaffected by Δ*rny* (Figs. 4, S3, and S5). Together, these data confirm that the cellular amounts of a relatively small number of sRNAs are changed in the Δ*rny* mutant.

**Figure 3:**
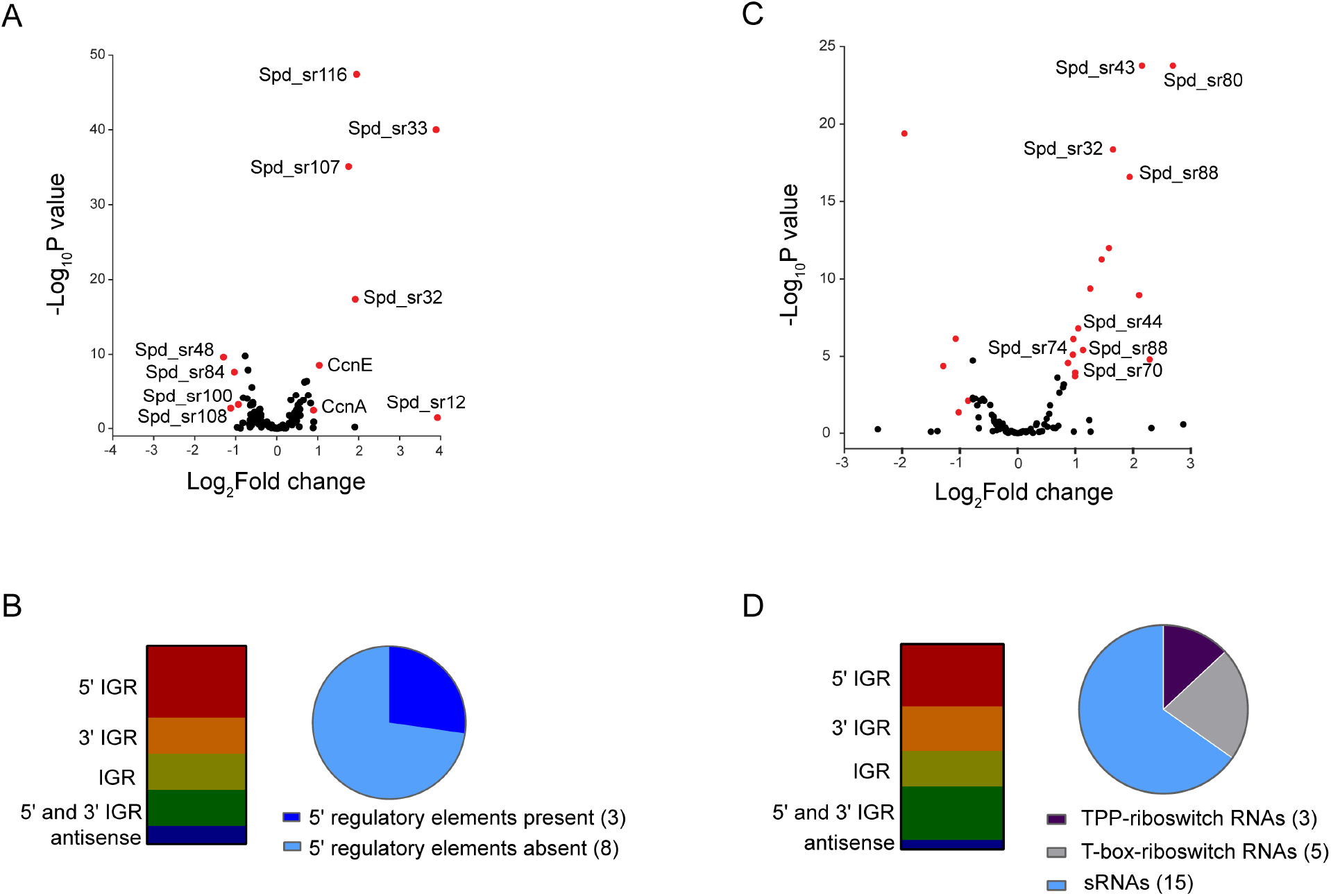
Impact of RNase Y and PNPase on sRNA transcriptome of *S. pneumoniae* D39. (A, C) Volcano plot showing genome-wide changes in sRNA transcript levels in a Δ*rny* mutant (A) or Δ*pnp* mutant (C) relative to the D39 parent strain. RNA was extracted from exponentially growing cultures of the WT D39 parent (IU1781) and isogenic mutants, Δ*rny* mutant (NRD10092) or Δ*pnp* mutant (IU4883), in triplicate and analyzed by sRNA-seq analysis as described in *Materials and Methods*. Red and blue dots represent genes with relative transcript changes >1.8- fold as the cut-off (Log_2_fold change =0.85), with an adjusted P-value cut-off <0.05. Relative transcript level changes of genes below the cut-off values are considered to be insignificant and are color coded in black. The X-axis represents gene-fold changes, and the Y-axis represents corresponding P-values plotted on a logarithmic scale. sRNAs that were significantly up-regulated or down-regulated in the Δ*rny* mutant or Δ*pnp* mutant compared to the parent are listed in Table 3 and Table 4, respectively. (B, D) Distribution of sRNAs that were differentially regulated in a Δ*rny* mutant (B) or a Δ*pnp* (D) mutant compared to the parent in different genomic contexts as described previously (40). Pie-chart graphs indicate the percent distribution of the sRNAs based on the presence or absence of 5’-cis-regulatory elements in their sequence. IGR, intergenic region.

**Figure 4:**
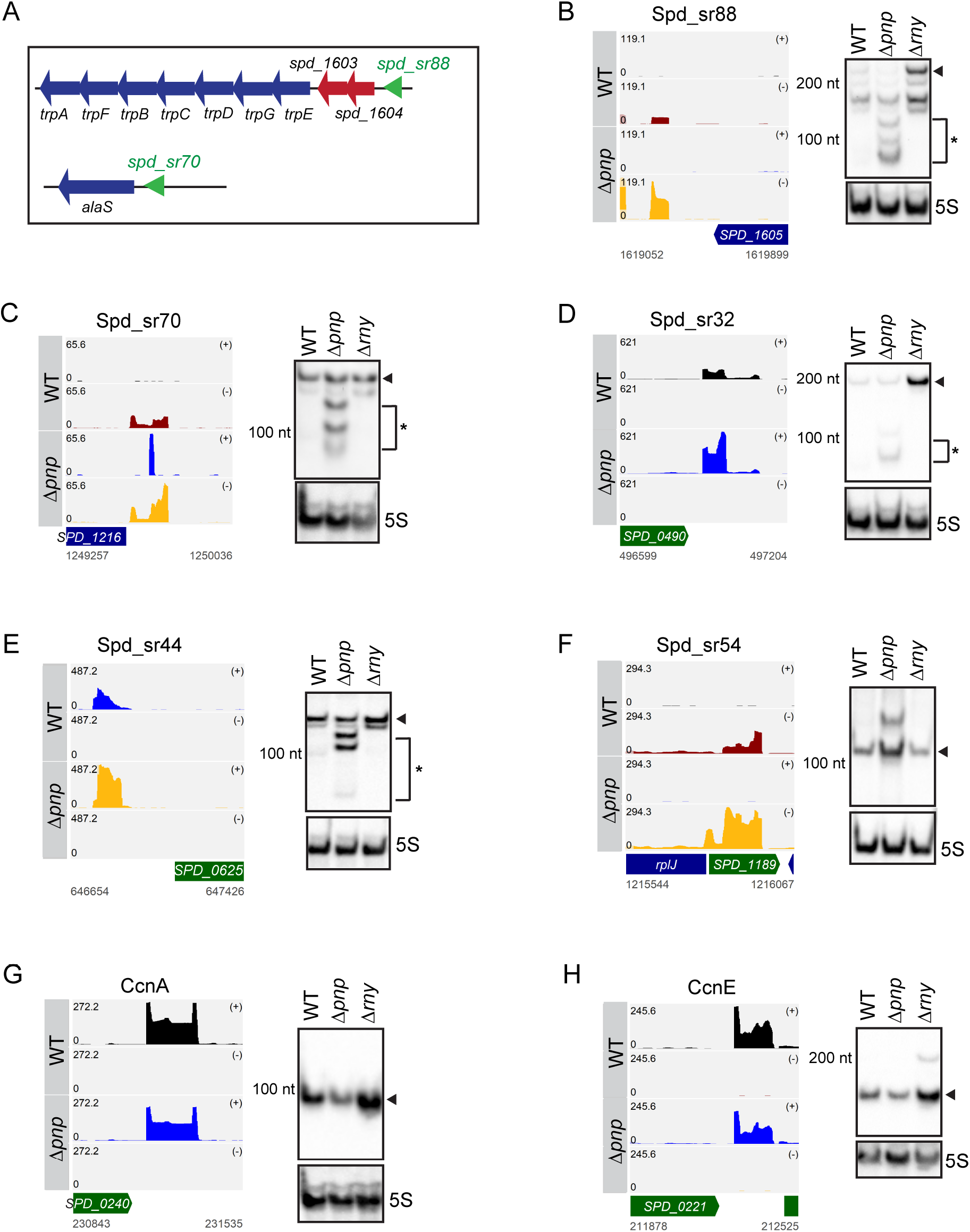
PNPase plays an important role in the decay and processing of riboswitch RNAs in *S. pneumoniae* D39. (A) Genetic context of two T-box riboswitches, Spd-sr88 and Spd-sr70, in *S. pneumoniae* D39. (B-H) Read coverage maps of a subset of sRNAs and their flanking regions that were differentially regulated in a Δ*pnp* mutant (IU4883) compared to the WT parent (IU1781) in sRNA-seq. Coverage represents depth per million reads of paired-end sRNA fragments and was averaged between normalized replicates (see Materials and Methods). In each coverage graph, ORFs encoded on the plus or minus strand are color-coded in green or blue, respectively. Northern blots detecting the sRNAs are presented alongside the read coverage maps. Black triangles and asterisks (*) indicate the full-length sRNA transcripts and sRNA decay products, respectively. Corresponding coverage maps for the sRNAs presented in panels B-H, in a Δ*rny* mutant (NRD10092) compared to the WT parent (IU1781) are presented in Figure S4. Quantification of signal intensity for each full-length sRNA normalized to 5S rRNA amount is displayed in Fig S5, and the probes used are listed in Table S3.

**Table 3:**
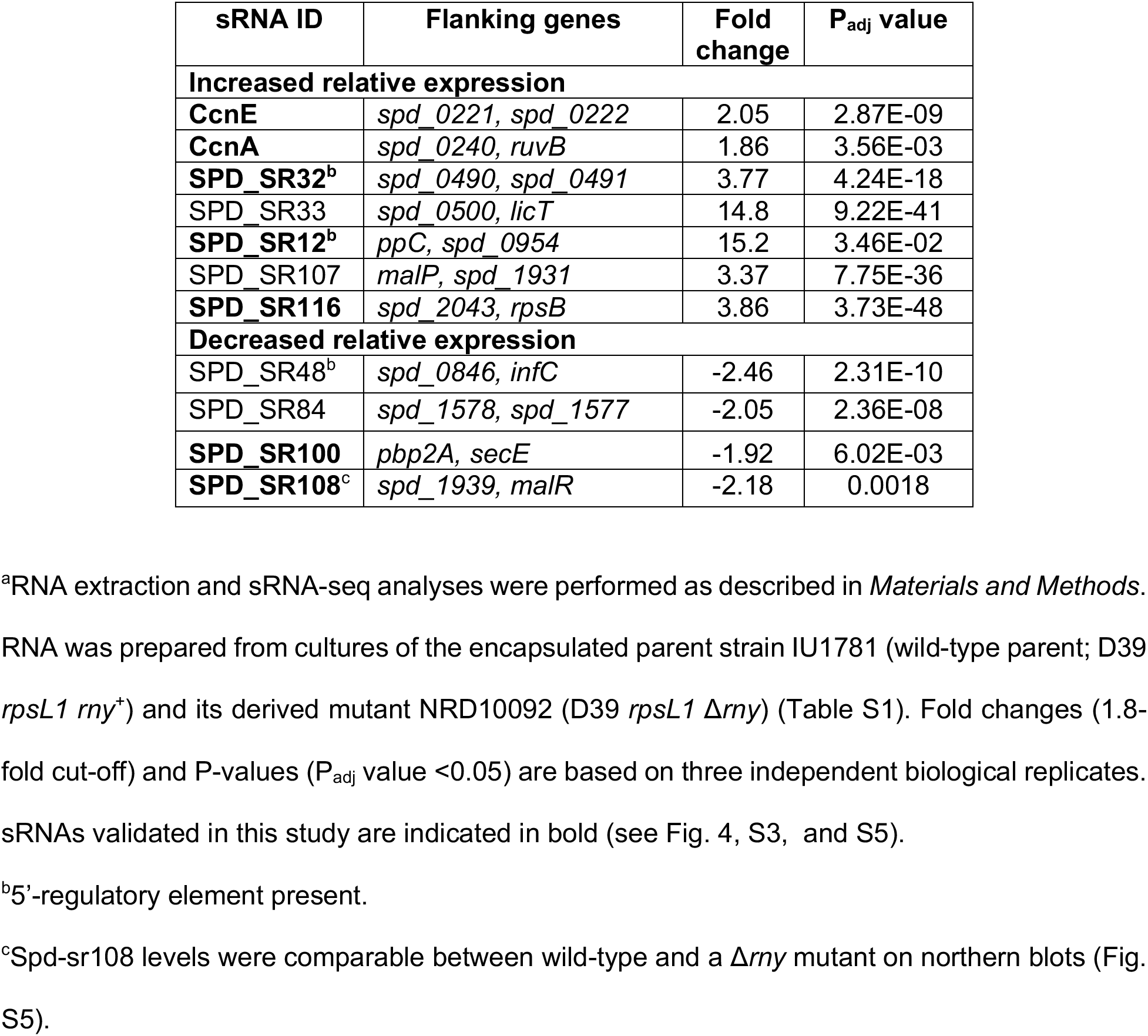
Relative sRNA transcript level changes in strain a Δ*rny* mutant compared to the *rny*^+^ parent strain during exponential growth in BHI broth^a^

In contrast to the Δ*rny* mutant, 21% of the pneumococcal sRNA transcriptome is significantly altered in the Δ*pnp* mutant. Twenty-three sRNAs exhibit >1.8-fold difference in relative expression in the Δ*pnp* mutant (Table 4; Fig. 3C), where 17 and 6 sRNAs are significantly up- and down- regulated, respectively. Notably, approximately half of the sRNAs up-regulated in a Δ*pnp* mutant relative to the WT strain are riboswitch RNAs. Spd-sr32, Spd-sr70, Spd-sr74, Spd-sr80, and Spd- sr88 are characterized by the presence of a T-box riboswitch, while Spd-sr43, Spd-sr44, and Spd- sr114 contain a TPP riboswitch element (Table 4; Figs. 3C and 3D). The two riboswitch RNAs Spd-sr44 and Spd-sr88 are particularly interesting, because Tn-seq screens with the serotype 4 strain (TIGR4) of *S. pneumoniae* indicated that the loss of *spd-sr44* or *spd-sr88* results in reduced pneumococcal fitness in murine blood and lung infection, respectively (44). 5’-intergenic and 3’- intergenic sRNAs are among the overrepresented category of sRNAs that show differential regulation in Δ*pnp* compared to the WT strain (Fig. 3D). Finally, we validated the expression of a total of 14 out of 23 sRNAs that were significantly differentially expressed in the Δ*pnp* mutant relative to the WT strain (Fig. 4, Fig. S3, and S5). Taken together, these data suggest that both RNase Y and PNPase play an important role in regulating the relative amounts of different sets of pneumococcal regulatory RNAs.

**Table 4:**
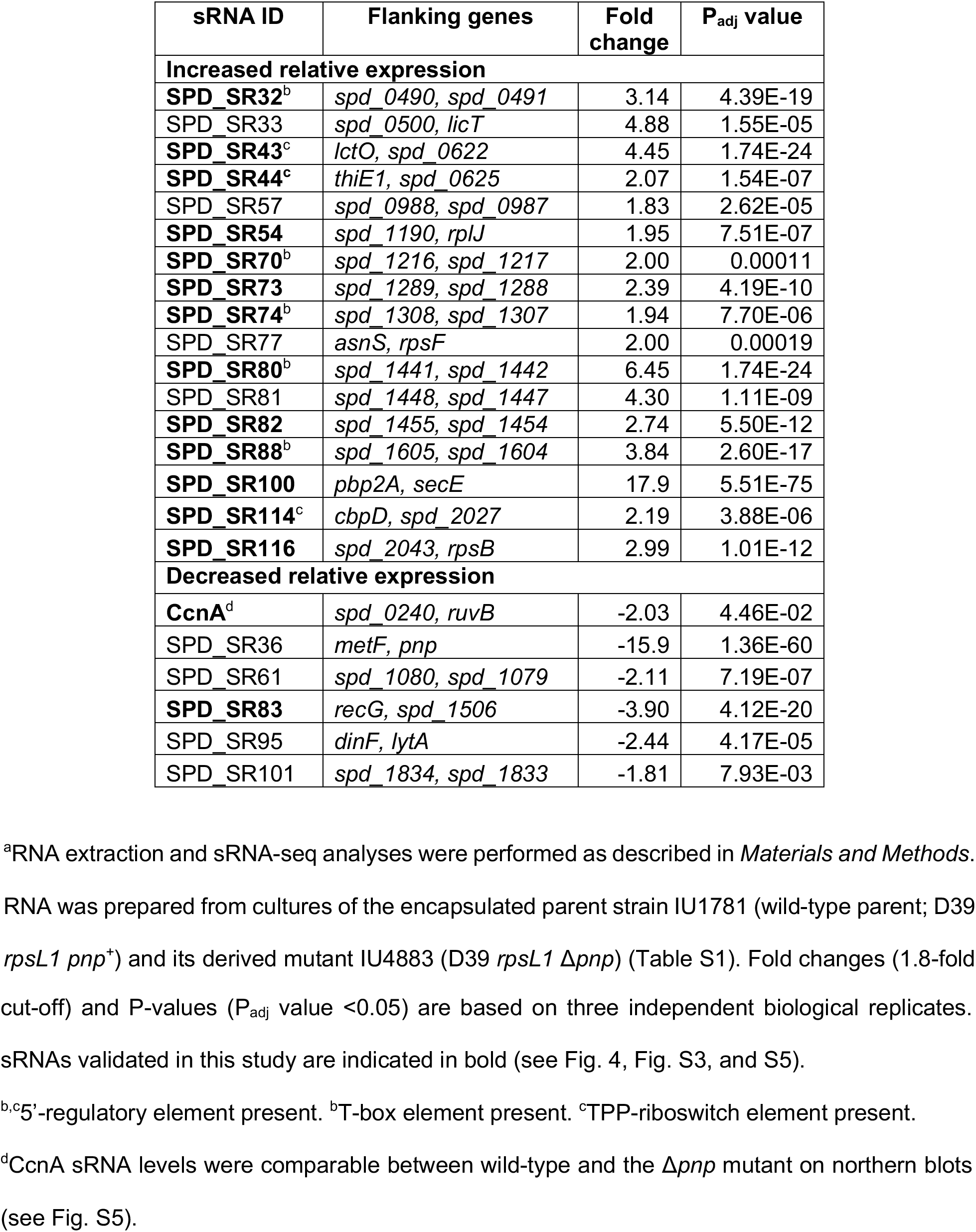
Relative sRNA transcript level changes in a Δ*pnp* mutant compared to the *pnp*^+^ parent strain during exponential growth in BHI broth^a^

| sRNA ID | Flanking genes | Fold change | P<sub>adj</sub> value |
| --- | --- | --- | --- |
| **Increased relative expression** |  |  |  |
| **SPD_SR32<sup>b</sup>** | *spd_0490, spd_0491* | 3.14 | 4.39E-19 |
| SPD_SR33 | *spd_0500, licT* | 4.88 | 1.55E-05 |
| **SPD_SR43<sup>c</sup>** | *lctO, spd_0622* | 4.45 | 1.74E-24 |
| **SPD_SR44<sup>c</sup>** | *thiE1, spd_0625* | 2.07 | 1.54E-07 |
| SPD_SR57 | *spd_0988, spd_0987* | 1.83 | 2.62E-05 |
| **SPD_SR54** | *spd_1190, rplJ* | 1.95 | 7.51E-07 |
| **SPD_SR70<sup>b</sup>** | *spd_1216, spd_1217* | 2.00 | 0.00011 |
| **SPD_SR73** | *spd_1289, spd_1288* | 2.39 | 4.19E-10 |
| **SPD_SR74<sup>b</sup>** | *spd_1308, spd_1307* | 1.94 | 7.70E-06 |
| SPD_SR77 | *asnS, rpsF* | 2.00 | 0.00019 |
| **SPD_SR80<sup>b</sup>** | *spd_1441, spd_1442* | 6.45 | 1.74E-24 |
| SPD_SR81 | *spd_1448, spd_1447* | 4.30 | 1.11E-09 |
| **SPD_SR82** | *spd_1455, spd_1454* | 2.74 | 5.50E-12 |
| **SPD_SR88<sup>b</sup>** | *spd_1605, spd_1604* | 3.84 | 2.60E-17 |
| **SPD_SR100** | *pbp2A, secE* | 17.9 | 5.51E-75 |
| **SPD_SR114<sup>c</sup>** | *cbpD, spd_2027* | 2.19 | 3.88E-06 |
| **SPD_SR116** | *spd_2043, rpsB* | 2.99 | 1.01E-12 |
| **Decreased relative expression** |  |  |  |
| **CcnA<sup>d</sup>** | *spd_0240, ruvB* | -2.03 | 4.46E-02 |
| SPD_SR36 | *metF, pnp* | -15.9 | 1.36E-60 |
| SPD_SR61 | *spd_1080, spd_1079* | -2.11 | 7.19E-07 |
| **SPD_SR83** | *recG, spd_1506* | -3.90 | 4.12E-20 |
| SPD_SR95 | *dinF, lytA* | -2.44 | 4.17E-05 |
| SPD_SR101 | *spd_1834, spd_1833* | -1.81 | 7.93E-03 |

<sup>a</sup>RNA extraction and sRNA-seq analyses were performed as described in *Materials and Methods*. RNA was prepared from cultures of the encapsulated parent strain IU1781 (wild-type parent; D39 *rpsL1 pnp<sup>+</sup>*) and its derived mutant IU4883 (D39 *rpsL1 Δpnp*) (Table S1). Fold changes (1.8-fold
cut-off) and P-values ( $P_{\text{adj}}$ value $<0.05$ ) are based on three independent biological replicates.
sRNAs validated in this study are indicated in bold (see Fig. 4, Fig. S3, and S5).
<sup>b,c</sup>5'-regulatory element present. <sup>b</sup>T-box element present. <sup>c</sup>TPP-riboswitch element present.
<sup>d</sup>CcnA sRNA levels were comparable between wild-type and the $\Delta pnp$ mutant on northern blots (see Fig. S5).

### PNPase and RNase Y play an important role in riboswitch RNA decay and processing

T-box containing riboswitch RNAs that are up-regulated in the Δ*pnp* mutant include Spd-sr88 and Spd-sr70, which are located within the 5’-UTRs of the *trp* operon (encoding enzymes involved in tryptophan biosynthesis) and *alaS* (encoding alanyl-tRNA synthetase), respectively (Table 4; Fig. 4A). Northern blotting confirmed increases in *spd-sr88* and *spd-sr70* in the Δ*pnp* mutant compared to the WT strain determined by RNA-seq analysis (Table 4) and showed accumulations of decay products of these sRNAs (Fig. 4B and 4C). Concurrently, both relative transcript amount of *alaS* and the entire *trp* operon, including upstream gene *spd_1604*, are decreased by ≈2-fold and ≈2-4-fold, respectively, in the Δ*pnp* mutant in mRNA-seq analysis (Table 2; Fig. 3C and 3D). Based on these observations, we further tested the expression profiles of six other TPP or T-box riboswitch RNAs that also showed increased relative expression in the Δ*pnp* mutant (Table 4). We observed a similar pattern of accumulation of decay intermediates for Spd-sr32, Spd-sr43, Spd-sr44, Spd-sr74, Spd-sr80 and Spd-sr114 in the Δ*pnp* mutant, but not in the WT or Δ*rny* strain (Fig. 4D, 4E, and Fig. S3). In addition, we noticed that the full-length transcripts for all six of these riboswitch RNAs are more abundant in a Δ*rny* mutant than that in a WT strain (Fig. 4B, 4D, Fig. S3, and S4). While Spd-sr32 was identified by sRNA-seq analysis to be significantly up-regulated by ≈4-fold in the Δ*rny* mutant compared to WT (Table 3), the increased relative transcript steady-state levels of the other riboswitch RNAs (Spd-sr43, Spd-Sr44, Spd-sr74, Spd-sr88, and Spd- sr114) in a Δ*rny* mutant were not detected by this transcriptomic-based approach, but were detected independently by northern-blot analysis (Fig. S3). Taken together, we conclude that both RNase Y and PNPase jointly function in the processing and decay of riboswitch regulatory RNAs.

### RNase Y regulates Ccn sRNA stability and function

After validating by northern blotting, the ≈3-fold increases in relative steady-state levels of CcnA and CcnE in the Δ*rny* mutant (Figs. 4G, 4H, 5A, 5B, 5D, 5E, and S6) in accordance with our sRNA-seq data (Table 3), we sought to further define the mechanism by which RNase Y regulates of the abundance of the Ccn sRNAs in *S. pneumoniae* D39. Therefore, we measured the stability of CcnA and CcnE in exponentially growing cultures of a Δ*rny* mutant or a WT strain after blocking transcription initiation by adding rifampicin (Fig. 5C and 5F). The relative half-life of CcnA increased by approximately 3-fold in Δ*rny* compared to WT (t_1/2_= 52.2 min versus t_1/2_= 17.6 min (Fig. 5C)), while that of CcnE increased by ≈2-fold (t_1/2_= 28.4 min versus t_1/2_= 15.8 min (Fig. 5F)) (Table S4). These findings prompted us to test the impact of RNase Y on the stability and the corresponding steady-state levels of the remaining three Ccn sRNAs. The relative transcript levels for CcnB, CcnC and CcnD, were similarly up-regulated by ≈2-fold in the Δ*rny* mutant compared to WT (Fig. 5G). Consistent with these increased amounts, CcnB and CcnC were significantly stabilized in the Δ*rny* mutant (Figs. 5H and 5I; Table S4). We were unable to accurately determine the relative stability of CcnD, because it was extremely unstable in the WT strain following rifampicin addition (data not shown).

**Figure 5:**
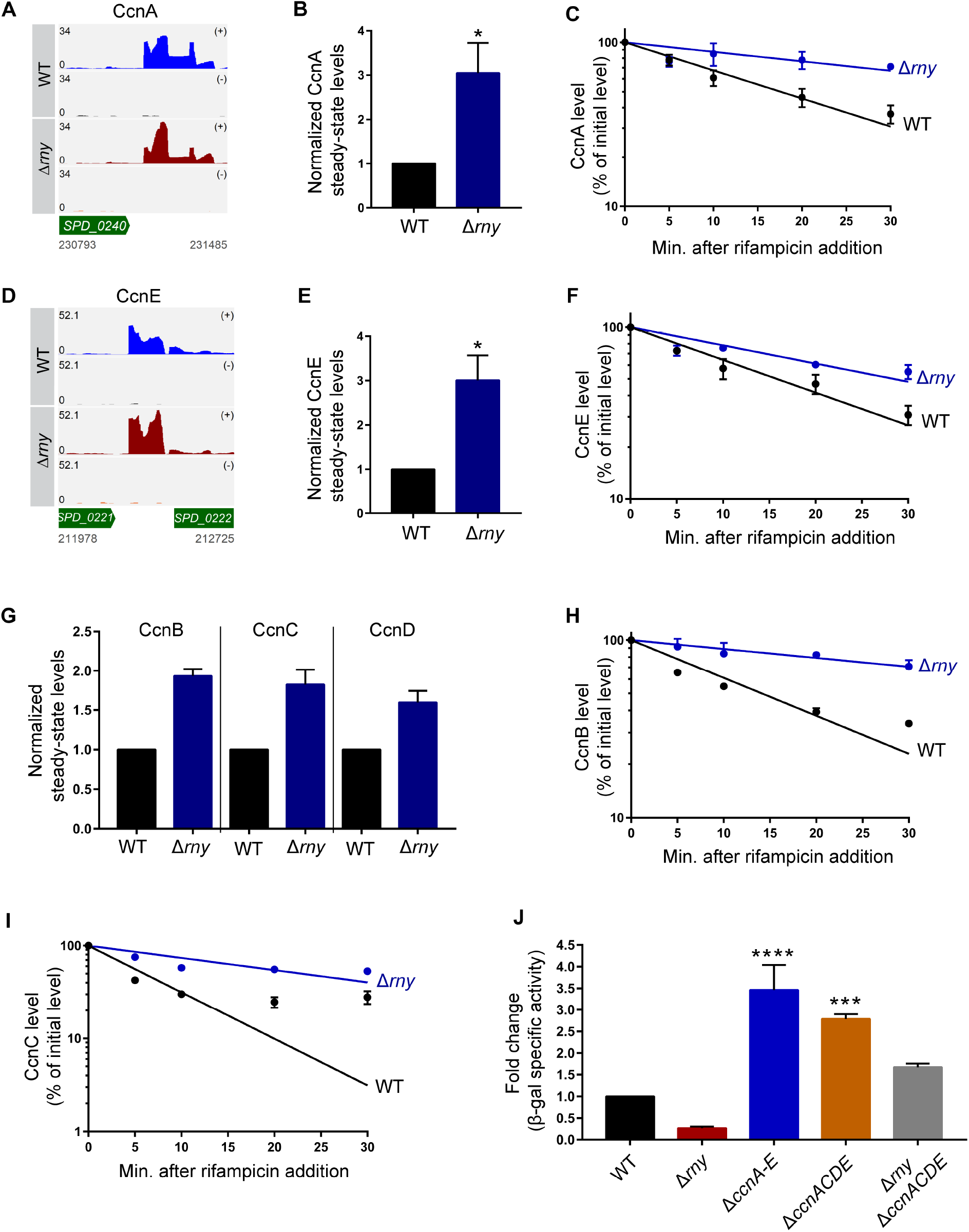
RNase Y regulates Ccn sRNA stability and function in *S. pneumoniae* D39. (A, D) Read coverage maps of CcnA and CcnE in a Δ*rny* mutant (NRD10092) compared to the WT parent (IU1781). Track labels corresponding to read coverage maps are described in the legend to Fig. 4. (B, E, G) CcnA, CcnE, CcnB, CcnC, and CcnD steady-state levels were determined on northern blots following extraction of RNA from exponentially growing cultures of a Δ*rny* mutant (NRD10092) and a WT parent strain (WT; IU1781) as described in Materials and Methods. Signal intensities in the northern blots were quantified and normalized to 5S RNA amount. (C, F, H, I) RNA amount time course experiment to determine the intrinsic stability of CcnA, CcnE, CcnB, and CcnC in a Δ*rny* mutant (NRD10092) and the WT strain (IU1781) after treatment with rifampicin to stop transcription, as described in Material and Methods. Semi-log sRNA decay curves were generated by fitting the normalized signal intensities determined on northern blots for each time-point sample. Points and error bars in the curves (where not visible, error bars are smaller than the symbol) represent the means (±SEM) of at least three independent experiments. sRNA half-life measurements corresponding to RNA stability curves are listed in Table S4. (J) β- galactosidase assay to determine the impact of RNase Y on Ccn sRNA-mediated *comC* translational regulation. Expression of the *comC*’-’*lacZ* translational fusion was monitored by β- galactosidase assays of samples removed from exponentially growing cultures of unencapsulated D39 parent strain (NRD10041) and isogenic mutant strains, Δ*cps* Δ*rny comC*’-’*lacZ* (NRD10113), Δ*cps* Δ*ccnA-E comC*’-’*lacZ* (NRD10187), Δ*cps* Δ*ccnACDE comC*’-’*lacZ* (NRD10054), and Δ*cps* Δ*rny* Δ*ccnACDE comC*’-’*lacZ* (NRD10120). Data and error bars represent the means (± SEM) of at least three independent experiments and asterisks (*), (**), (***) and ns indicate P<0.05, P<0.01, P<0.001, and not significantly different, respectively.

Finally, we investigated the role of RNase Y in Ccn-mediated *comC* target regulation. To this end, we constructed a translational fusion in which the 5’ untranslated region and the first twelve codons of *comC* are fused in-frame with the truncated *E. coli* β-galactosidase gene *lacZ*. The *comC*’-’*lacZ* translational fusion, driven from the constitutive vegetative promoter *vegT* (derived from the *vegII* promoter of *Bacillus subtilis*; see Table S1 and S2), was integrated in the chromosomal *bgaA* locus in strain D39 (thereby, knocking out pneumococcal β-galactosidase). Consistent with previous reports, deletion of all 5 Ccn sRNAs (Δ*ccnA-E*) relieved ComC translational repression and led to increased relative expression of β-galacosidase specific activity (≈3.5-fold) from *comC*’-’*lacZ* (Fig. 5J). Conversely, Δ*rny* led to decreased (≈3.8-fold) relative β-galactosidase specific activity from *comC*’-’*lacZ* (Fig. 5J), consistent with increased stabilization of Ccn sRNAs (Fig. 4) and increased translational repression of ComC. To further test this idea, we attempted to measure *comC*’-’*lacZ* expression in a Δ*rny* Δ*ccnA-E* mutant. Unexpectedly, the Δ*rny* Δ*ccnA-E* mutant exhibited a synthetic phenotype with severely impaired growth and low growth yield compared to WT (data not shown). By contrast, a Δ*rny* Δ*ccnACDE* mutant did not exhibit a strong synthetic phenotype (data not shown). Relative expression of *comC*’-’*lacZ* is less elevated in the Δ*ccnACDE* than the Δ*ccnA-E* mutant and is reduced further in the Δ*rny* Δ*ccnACDE* to near the WT level (Fig. 5J), consistent with stabilization of remaining CcnB in the Δ*rny* background. Altogether, these results indicate that RNase Y-mediated regulation of Ccn sRNA stability has a consequential impact on Ccn-mediated target regulation in *S. pneumoniae* D39.

## DISCUSSION

This paper is the first report of the global roles of two highly conserved Gram-positive RNases, RNase Y and PNPase, in the human pathogen *S. pneumoniae*. The loss of RNase Y significantly impacts gene expression by affecting ≈10% of the pneumococcal transcriptome and thereby causing pleiotropic phenotypes (Fig. 1A, 1C, 1D, 2A, 2B, 3A, and 3B; Tables 1 and 3). In contrast, PNPase exerts a relatively smaller impact on the transcriptome compared to RNase Y, but interestingly regulates the expression of specific transcripts previously implicated in pneumococcal virulence control (Tables 2 and 4 and Fig. 2C, 2D, 3C and 3D). Accordingly, the loss of PNPase severely attenuates *S. pneumoniae* virulence *in vivo* (Fig. 1E and S2). This study also uncovered that both RNase Y and PNPase work in concert to regulate the processing and decay of several regulatory RNAs; in particular, those characterized by the presence of 5’-cis-acting regulatory elements (Fig. 3B, 3D, 4, and S3; Tables 2 and 4). In addition, RNase Y stabilizes the conserved pneumococcal trans-acting sRNAs CcnA-E, further impacting Ccn- mediated target gene regulation (Fig. 5).

### RNase Y is a pleiotropic regulator in *S. pneumoniae* D39

Deletion of *rny* in *S. pneumoniae* leads to a ≈2-fold increase in doubling time *in vitro* (Fig. 1A and Table S2), similar to prior observations with *B. subtilis* and *C. perfringens* (25, 27) and interferes with pneumococcal cell division (Fig. 1C). We identified several important pneumococcal cell wall and division regulators, including *mapZ* (encoding a mid-cell-anchor protein), *cozE* (encoding a coordinator of zonal division), and *gpsB* (encoding a regulator that balances septal and elongation peptidoglycan synthesis) as up-regulated significantly in a Δ*rny* mutant (Fig. 2B; Table 1). In *S. pneumoniae*, MapZ guides tubulin-like FtsZ protein from mid-cell rings of dividing cells to the equators of new daughter dells (45, 46), whereas, GpsB and CozE are major peptidoglycan (PG) biosynthesis regulators that play distinct but crucial roles at the midcell to maintain the normal ovococcus shape of pneumococcus by modulating the activities of different penicillin-binding proteins (PBPs), which catalyze peptide cross-link formation in peptidoglycan (47–49). Accordingly, Δ*mapZ* mutants exhibit a variety of abnormal cell shapes and sizes, decreased cell viability, increased doubling time, and aberrant FtsZ movement (45, 46) while cells depleted for *gpsB* or *cozE* form elongated cells that are unable to divide or form chains that round up and lyse, respectively (47, 49, 50).

Several transcripts under the control of the essential TCS WalRK and TCSs LiaFSR and CiaRH are impacted in the Δ*rny* mutant (Table 1), again consistent with cell wall and surface stress in cells lacking RNase Y as numerous proteins in these regulons are known to impact cell morphology and chaining (51–55). In this regard, the defects in cell shape and morphology observed for *B. subtilis* Δ*rny* mutants were attributed to the up-regulation of several PG biosynthesis genes, including *rodA* (27). It remains to be determined what cell wall stress is caused by absence of pneumococcal RNase Y and whether induction of certain proteins in these multiple surface-stress TCS regulons can account for the defects in growth and morphology of the *S. pneumoniae* Δ*rny* mutant.

Besides responding to cell-wall stress, the CiaRH TCS has been implicated in pneumococcal biofilm formation (56), competence (57), and virulence (58). In particular, the 5 five conserved pneumococcal base-pairing sRNAs (CcnA-E) negatively regulate translation of *comC*, which encodes the competence stimulatory peptide (42, 43, 44). We show here that RNase Y functions as a critical regulator of Ccn sRNA stability and impacts Ccn-mediated negative regulation of competence development in *S. pneumoniae* (Fig. 5). Interestingly, the recent Grad-seq analysis indicated possible stable RNA-protein complexes between the 3’-to-5’ exonuclease YhaM/Cbf1 and the Ccn sRNAs that were confirmed in pull-down experiments with Ccn sRNAs as bait in *S. pneumoniae* TIGR4 strain. In addition, CcnA-E pulled down several degradosome components, including RNase J1/J2 and PNPase (23). The Gram-positive specific Cbf1 exonuclease has been implicated in trimming single-stranded RNA tails at the 3’-ends of Rho-independent terminated transcripts (16, 17, 23), thereby, preventing decay by other exoribonucleases, such as PNPase and RNase R that require an unstructured tail of 7-10 nt for binding (17, 59). Although data presented here suggests that the Ccn sRNAs are targeted by RNase Y (Fig. 5), RNase Y was not identified as a strong Ccn-sRNA interactor by Grad-seq (23), perhaps indicating complex dissociation during gradient centrifugation. Future experiments will determine whether Ccn sRNAs are direct substrates of RNase Y and whether Cbf1-mediated 3’ trimming impacts Ccn sRNA decay by RNase Y. Moreover, results in this paper raise the important question of whether RNase Y functions similarly to RNase E in mediating decay of trans-acting sRNAs that form sRNA-mRNA base-pairs in *S. pneumoniae* and other Gram-positive bacteria.

Finally, lack of RNase Y affected the steady-state transcript levels of numerous key metabolic operons and known pneumococcal colonization and virulence factors, including *pavB* (fibronectin-binding protein; host interaction) (60); *clpL* (adaptor protein for ClpP protease) (61); CiaRH TCS regulon members (*licC, licB, and licA* (choline metabolism)) (62), LiaFSR TCS regulon members (*dnaK* and *dnaJ* (protein chaperones)) (63); WalRK TCS regulon members (*lytB* (glucosaminidase)) *spd_0104* (LysM-protein) (35); and PnpRS TCS regulon members (phosphate uptake) (35, 64) (see Table 1). Thus, loss of RNase Y clearly exerts a global impact on the pneumococcal transcriptome that broadly affects physiology, growth, and virulence.

### PNPase is a key regulator of *S. pneumoniae* D39 virulence

In contrast to the highly pleiotropic effects caused by the absence of RNase Y, the lack of PNPase minimally affects growth or morphology *in vitro*, but remarkably, causes strong attenuation *in vivo* (Fig. 1A, 1B, 1C and 1E). The lack of phenotypes of the Δ*pnp* mutant *in vitro* may suggest that pneumococcal 3’- 5’ exoribonuclease RNase R can functionally bypass PNPase in certain experimental conditions. Notably, 10 out of 20 protein-coding transcripts that were either up- (*ribU* (≈4-fold), *fruR* (≈2-fold), *galE-2* (≈2-fold)) or down- (*trpACDGE* (≈2.5-4-fold)) regulated in the Δ*pnp* mutant included metabolic genes implicated in nasopharyngeal colonization and/or lung infection in a mouse model (35) (Fig. 2C and 2D; Table 2). The relative level of the full-length transcript of the T-box riboswitch Spd-sr88 located within the 5’-UTR of the *trp* operon (Fig. 4A) also decreased by ≈2- fold in the Δ*pnp* mutant, with concomitant accumulation of *spd-sr88* derived-decay intermediates (Fig. 4B; Table 4). These decay products are likely generated by RNase Y cleavage, since the relative full-length Spd-sr88 transcript levels increase by ≈11-fold in a Δ*rny* mutant (Fig. 4B and S5).

We do not yet know how PNPase positively regulates *trp* operon in *S. pneumoniae* but in general, *trp* operon regulation is important and complex in different bacteria and often involves RNA-based post-transcriptional mechanisms (65). For example, in *B. subtilis* under tryptophan replete conditions, *trp* expression is repressed as a consequence of TRAP regulator protein mediated transcription termination of the *trp* leader which is subsequently degraded by RNase Y and/or J1 and PNPase. (66). In *E. coli*, tryptophan synthesis is regulated by a classical transcription attenuation mechanism, where under tryptophan replete conditions the upstream *trpL* leader peptide (TrpL) is translated efficiently allowing formation of a terminator stem-loop that stops transcription before the downstream *trp* genes (67). Recently, the terminated *trpL* RNA generated by this attenuation mechanism was shown to function in *Sinorhizobium meliloti* as a base-pairing sRNA to destabilize several transcripts, including that of the *trp* biosynthesis genes (68). Likewise, previous studies in important Gram-positive pathogens, including *Listeria monocytogenes* and *Enterococcus faecalis*, show that terminated riboswitches are not just “junk RNA,” but function as mRNA- or protein-binding regulatory RNAs (69, 70). *S. pneumoniae* does not possess obvious homologs of TRAP or TrpL, but its *trp* operon instead contains the T-box(tRNA-sensing structure) riboswitch Spd-sr88. Whether the RNA decay products derived from *spd-sr88* (Fig. 4B) function as regulatory RNAs to destabilize the *trp* operon transcript in a Δ*pnp* mutant awaits further investigation. These combined results show that PNPase controls the transcript amounts of numerous genes required for pneumococcal pathogenesis, including the *trp* operon and the riboswitches Spd-sr88 and Spd-sr44 (Fig. 4B, 4E, S5; Table 2 and 4) (35, 44), supporting the notion that PNPase is a key regulator of *S. pneumoniae* pathogenesis.

### RNase Y and PNPase play roles in sRNA processing and decay in *S. pneumoniae* D39

Riboswitch turnover is important for recycling of the ligands to which they respond, and a role for RNase Y in this process has previously been reported in *B. subtilis* (11, 71) and *S. aureus* (21). Here, we show that pneumococcal RNase Y mediates the initial endo-ribonucleolytic cleavage of 5’-cis-acting regulatory elements, which are subsequently degraded by PNPase. Likewise, in a Δ*pnp* mutant, we found that decay intermediates of eight riboswitch RNAs accumulated, while their corresponding full-length transcripts increased in abundance in the absence of RNase Y (Figs. 4, S3, and S5; Tables 3 and 4). These observations are consistent with a recent study in *S. pyogenes* showing that the coordinated actions of RNase Y and PNPase play a crucial role in the decay of riboswitches (20). In addition, our data indicate that RNase Y likely generates some sRNAs by cleaving larger transcripts, as observed for Spd-sr88 and Spd-sr116 (Fig. 4, Fig. S3, and S5). We conclude that RNase Y and PNPase work in tandem to degrade pneumococcal cis-acting regulatory RNAs, while RNase Y also plays an important role in sRNA processing and maturation. Whether RNase Y and PNPase interact together in a degradosome-like complex to impact regulatory RNA levels in *S. pneumoniae* will be resolved in future experiments.

## MATERIALS AND METHODS

### Bacterial strains and growth conditions

Bacterial strains used in this study were derived from encapsulated *S. pneumoniae* serotype 2 strain D39W and are listed in Table S1. Strains were grown on plates containing trypticase soy agar II (modified; Becton-Dickinson [BD]) and 5% (vol/vol) defibrinated sheep blood (TSAII BA) at 37°C in an atmosphere of 5% CO2. Liquid cultures were grown statically in BD brain heart infusion (BHI) broth at 37°C in an atmosphere of 5% CO_2_. Bacteria were inoculated into BHI broth from frozen cultures or single colonies. For overnight cultures, strains were first inoculated into 17-mm-diameter polystyrene plastic tubes containing 5 mL of BHI broth and then serially diluted by 100-fold into five tubes; these cultures were then grown for 10 to 16 h. Cultures with an OD_620_ = 0.1 to 0.4 were diluted to a starting OD_620_ between 0.002 and 0.005 in 5 mL of BHI broth in 16-mm glass tubes. Growth was monitored by measuring OD_620_ using a Spectronic 20 spectrophotometer. For antibiotic selections, TSAII BA plates and BHI cultures were supplemented with 250 µg/mL kanamycin or 150 µg/mL streptomycin.

### Construction and verification of mutants

Mutant strains were constructed by transformation of competent *S. pneumoniae* strains with linear PCR amplicons as described previously (72). DNA amplicons containing antibiotic resistance markers were synthesized by overlapping fusion PCR. *S. pneumoniae* cells were induced to competence by the addition of synthetic competence stimulatory peptide 1 (CSP-1; Anaspec, Inc.). Markerless deletions and replacements of target genes were constructed using the *kan*^R^-*rpsL*^+^ (Janus cassette) allele replacement method as described previously (73). In the first step, the Janus cassette was used to disrupt target genes in an *rpsL1* (Str^R^) strain background, and transformants were screened for kanamycin resistance and streptomycin sensitivity. In the second step, the Janus cassette was replaced by a PCR amplicon containing the desired mutation or replacement lacking antibiotic markers, and the resulting transformants were screened for streptomycin resistance and kanamycin sensitivity. Final transformants were isolated as single colonies three times on TSAII BA plates containing antibiotics listed in Table S1 and subsequently grown for storage in BHI containing the appropriate antibiotic. All constructs were confirmed by PCR amplification and sequencing.

### Microscopy

After cultures reached an OD_620_ ≈0.1- 0.2, 1 mL was removed and centrifuged at 16,000 × *g* for 2 min at room temperature. Pellets were suspended in 50 μL of BHI broth. Cells were examined using a Nikon E200 phase-contrast microscope, and images were captured using an Imaging Source DFK 1920 x 1200 pixel camera. A total of over 100 chains from each of two independent cultures of each strain were counted to determine distributions of numbers of cells per chain.

### RNA extraction

RNA for high throughput sequencing was prepared as described previously (74). Briefly, strains were grown in 30 mL of BHI starting at an OD_620_ = 0.002 in 50 mL conical tubes. RNA was extracted from exponentially growing cultures of IU3116 (wild-type parent; D39 *rpsL1* CEP:: *kanrpsL^+^*) and its derived isogenic mutants IU5498 (D39 *rpsL1* Δ*pnp CEP:: kanrpsL^+^*) and IU5504 (D39 *rpsL1* Δ*rny CEP:: kanrpsL^+^*) at OD_620_ ≈ 0.1 from matched batches of BHI broth for mRNA-seq analysis or from IU1781 (wild-type parent; D39 *rpsL1*) and its derived markerless mutants IU4883 (D39 *rpsL1* Δ*pnp*) and NRD10092 (D39 *rpsL1* Δ*rny*) at OD_620_ ≈ 0.15 for sRNA-seq analysis using the FastRNA Pro Blue Kit (MP Bio) according to the manufacturer’s guidelines. RNA extracted for mRNA-seq analysis was purified with a miRNeasy minikit (Qiagen), which included an on-column DNase I (Qiagen) treatment. For sRNA-seq analysis, RNA was alcohol precipitated following extraction and subsequently subjected to DNase treatment (Turbo DNase; Ambion) following the manufacturer’s protocol. Sample mixtures (total reaction volume of 50 µL) were incubated with Turbo DNase for 30 min at 37°C, and each reaction was stopped by addition of 150 µL of DEPC-treated water and 200 µL of neutral phenol–chloroform-isoamyl alcohol (Fisher). DNase-treated RNA samples were phenol extracted and alcohol precipitated. To isolate RNA for droplet-digital PCR, RNA was extracted from exponential growth phase cultures following the same procedure as described above for sRNA-seq analysis. The amount and purity of all RNA samples isolated were assessed by NanoDrop spectroscopy (Thermo Fisher). RNA integrity of the samples used for RNA-seq library preparation was further assessed using the Agilent 2100 BioAnalyzer (Aligent Technologies).

### Library preparation and mRNA-seq

cDNA libraries were prepared from total RNA by the University of Wisconsin-Madison Biotechnology Center as described previously (40). Briefly, total RNA was subjected to rRNA-depletion using RiboZero^TM^ rRNA Removal Kit (EpiCentre Inc., Madison, WI, USA). Double stranded cDNA synthesis was performed with rRNA-depleted mRNA using ScriptSeq^TM^ v2 RNA-Seq Library Preparation kit (EpiCentre Inc., Madison, WI, USA) in accordance with the manufacturer’s standard protocol. The amplified libraries were purified using Agencourt AMPure^®^ XP beads. Quality and quantity were assessed using an Agilent DNA 1000 chip (Agilent Technologies, Inc., Santa Clara, CA, USA) and Qubit® dsDNA HS assay kit (Invitrogen, Carlsbad, California, USA), respectively. Libraries were standardized to 2 μM and cluster generation was performed using standard Cluster kits (v3) and Illumina Cluster Station. Single-end 100 bp sequencing was performed using standard SBS chemistry (v3) on an Illumina HiSeq2000 sequencer. Images were analyzed using the standard Illumina pipeline, version 1.8.2.

### Library preparation and sRNA-seq

sRNA libraries were prepared from total RNA as described previously (40). Briefly, 5 µg of DNase-treated total RNA was subjected to ribosomal RNA removal (RiboZero^TM^ rRNA Removal for Gram-positive bacteria, Illumina). rRNA depleted samples were then subjected to RNA fragmentation using the Ambion RNA fragmentation kit (AM8740). Fragmented RNA was subjected to RNA 5’-polyphosphatase (Epicenter) treatment, which was performed to facilitate the 5’-adapter ligation step. Small RNA libraries were generated by Macrogen using the TruSeq® small RNA library kit (Illumina). 100 bp paired-end read sequencing was performed using an Illumina HiSeq2000 sequencer.

### RNA-seq analysis

Raw sequencing reads from mRNA-seq were quality and adapter trimmed using Trimmomatic version 0.17 (75) with a minimum length of 90, while those corresponding to sRNA-seq were preprocessed for alignment with Cutadapt. The trimmed reads were mapped on the *Streptococcus pneumoniae* D39 (RefSeq NC_008533) genome and D39 plasmid pDP1 sequence (RefSeq NC_005022) using Bowtie2 (76). mRNA-seq and sRNA-seq analysis were performed as described previously using DESeq2 (74). Genes were defined as differentially expressed if their P_adj_ (P-value adjusted for multiple testing) was <0.005. Primary data from mRNA-seq and sRNA-seq analyses was submitted to the NCBI Gene Expression Omnibus (GEO). The accession numbers for the sRNA-seq data corresponding to wild-type samples used for comparison of Δ*rny* and Δ*pnp* mutants are GSE148867 and GSE123437 respectively. The sRNA-seq and data corresponding to Δ*rny* and Δ*pnp* mutants, and the mRNA-seq data corresponding to all strains has been deposited to GEO under the accession number GSE173392.

### Droplet Digital PCR (ddPCR) analysis

One microgram of DNA-free RNA was reverse transcribed using random hexamers and Superscript III Reverse Transcriptase (RT) (Invitrogen) following the manufacturer’s protocol. For each sample, a no RT (NRT) control reaction was performed. cDNA samples were diluted 1:10, 1:10^2^, 1:10^3^, or 1:10^6^, and 2 µL of each diluted RT and NRT-PCR sample was added to a 22 µL reaction mixture containing 11 µL of QX200^TM^ ddPCR ^TM^ Evagreen Supermix (Bio-Rad) and 1.1 µL of each 2 µM ddPCR primers (Table S3). A single no template control (NTC) for each ddPCR primer pair used in this study was included. Droplet generation from each reaction mixture was achieved via the QX200 Automated Droplet Generator (Bio-Rad) and end-point PCR was performed using a thermal cycler following the instructions from the manufacturer. A QX200 Droplet Reader (Bio-Rad) was used to analyze droplets from each individual reaction mixture, where PCR-positive and PCR-negative droplets were counted to provide absolute quantification of the target transcript. Data analysis was performed with QuantaSoft software (Bio-Rad), and the concentration of each target is expressed as copies per µL. Reactions were performed using cDNA from at least three independent biological replicates, and transcript copies were normalized to 16S rRNA (internal control). Normalized transcript copy numbers were used to calculate fold changes of transcripts corresponding to target genes in different sets of mutants relative to the WT parent. Statistical analysis was performed using Student’s t-test in GraphPad Prism version 7.0.

### RNA stability assay

To determine RNA stabilities, cultures were grown in BHI to exponential phase (OD_620_ ≈ 0.15) as described above and a culture sample (T_0_) was collected. Rifampicin was added to inhibit transcription and additional samples were collected 5, 10, 20, and 30 min after rifampicin addition. All samples were subjected to hot phenol lysis method as described previously (77). Briefly, 700 µL of sample was added to a mixture containing 800 µL of acid phenol– chloroform-isoamyl alcohol, pH 4.3 (Fisher Scientific) and 100 µL of lysis buffer (320 mM sodium acetate (pH 4.6), 8% (wt/vol) SDS, and 16 mM EDTA) equilibrated to 65°C. Samples were mixed at 65°C for 5 min, and centrifuged for 30 min at 4°C to separate phases. The upper aqueous phase was extracted a second time with an equal volume of neutral phenol–chloroform-isoamyl alcohol, pH 6.7 (Fisher Scientific). RNA was ethanol-precipitated and resuspended in DEPC-treated water. RNA concentration was measured using a Nano Drop 2000 (ThermoFisher Scientific).

### Northern blot analysis

Two micrograms of each RNA sample was loaded on 10% polyacrylamide gels containing 7 M urea or loaded onto 10% Criterion TBE-urea precast gels (Bio-Rad) and electrophoresed at 85 V. RNA samples were transferred to a Zeta-Probe GT membrane (Bio-Rad) using a Trans-Blot SD semidry transfer apparatus (Bio-rad) following the manufacturer’s guidelines. Transferred RNA was UV-crosslinked and hybridized overnight with 100 ng/mL of 5′ biotinylated DNA probe (Table S3) in ULTRAhyb (Ambion) hybridization buffer at 42°C. Blots were developed using a BrightStar BioDetect kit protocol (Ambion), imaged with a ChemiDoc MP imager (Bio-Rad), and quantified using Image Lab software version 5.2.1 (Bio-Rad). Signal intensity corresponding to each sRNA was normalized to that of 5S rRNA, which served as internal loading control. Decay curves corresponding to RNA stability time course experiments were generated by using GraphPad Prism version 7.0.

### Mouse models of infection

All procedures were approved in advance by the Bloomington Institutional Animal Care and Use Committee (BIACUC) or UTHealth Animal Welfare Committee and were performed according to recommendations of the National Research Council. Experiments were performed as described in (73) with the following changes. Male ICR mice (21- 24 g; Harlan) were anaesthetized by inhaling 4% isoflurane (Butler animal Health Supply) for 8 min. In two independent experiments, a total of 8 mice were intranasally inoculated with each bacterial strain to be tested. Bacteria were grown exponentially in BHI broth in an atmosphere of 5% CO_2_ to OD_620_ ≈ 0.1. Ten milliliters of culture were centrifuged for 5 min at 14,500 x *g* and then suspended in 1 mL 1 X PBS to yield ≈ 10^7^ CFU mL^-1^. CFU counts were determined by serial dilution and plating. Fifty microliters of suspensions were administered intranasally as described previously (72). Mice were monitored visually at 4 to 8 h intervals, and moribund mice were euthanized by CO_2_ asphyxiation followed by cervical dislocation (IU-Bloomington), which was used as the time of death in statistical analyses. Alternatively, isoflurane-anesthetized moribund mice were euthanized by cardiac puncture-induced exsanguination followed by cervical dislocation (UTHealth). Kaplan-Meir survival curves and log-rank tests were generated using GraphPad Prism 7.0 software.

### β-Galactosidase assays

Strains containing the *comC*’-’*lacZ* translational fusion were grown in BHI broth to exponential phase (OD_620_ ≈ 0.15). Samples were taken from each culture and assayed for β-galactosidase activity as described by Miller (78) with slight modifications. Briefly, 1 mL of culture was removed and centrifuged at 16,000 × *g* for 2 min at 4°C. Pellets were resuspended in 1 mL of Z-buffer containing 2-beta-mecaptoethanol at a final concentration of 0.27%. Each sample mixture was lysed by subsequent incubation at 37°C for 10 min following the addition of 10 μL of 5% (vol/vol) Triton. 100 μL of lysed culture samples were then assayed for β-galactosidase specific activity as described previously.

## SUPPLEMENTAL MATERIALS

Supplemental Materials are available for this article.

## ACKNOWLEDGEMENTS

We thank Doug Rusch (Indiana University Bloomington) for assistance with Illumina mRNA-Seq analyses. This work was supported by the McGovern Medical Startup funds and NIGMS grant RO1GM121368 (to D.S., J.F., K.C., and N.R.D.) and NIGMS grant R35GM131767 (to M.E.W). Study designed by D.S., M.E.W., and N.R.D.; experiments carried out by D.S., J.F., K.C., and N.R.D.; analysis done by D.S., J.F., and N.R.D.; and paper written by D.S., M.E.W., and N.R.D.

