## SUPPLEMENTAL MATERIAL for "Pivotal Roles for Ribonucleases in *Streptococcus pneumoniae* Pathogenesis"

**Running Title:** PNPase and RNase Y functions in *Streptococcus pneumoniae*

**Keywords:** RNase Y; polynucleotide phosphorylase; small RNAs; post-transcriptional regulation

### TABLE OF CONTENTS

#### SUPPLEMENTAL TABLES

**TABLE S1.** *S. pneumoniae* strains and primers used in this study

**TABLE S2.** Growth characteristics of Δ*rny* mutants used in this study

**TABLE S3.** Droplet-Digital PCR primers and oligonucleotide probes used in this study

**TABLE S4**. Half-life calculations

#### SUPPLEMENTAL FIGURES

**FIGURE S1.** Growth characteristics of Δ*rny* and Δ*pnp* mutants in different batches of BH both at optimal (37°C) and lower (32°C) temperatures.

**FIGURE S2.** Survival curve analysis showing disease progression of Δ*pnp* mutant compared to a wild-type parent in an invasive model of pneumonia.

**FIGURE S3**. Northern blot validations for sRNA-seq analysis.

**FIGURE S4.** Read coverage maps for sRNAs shown in Figure 4 in a Δ*rny* mutant compared to the wild-type parent.

**FIGURE S5.** Corresponding northern blot quantifications for Figure 4 and Figure S3.

**FIGURE S6.** Northern blots corresponding to data presented in Figure 5.

#### SUPPLEMENTAL REFERENCE

**Table S1.** *S. pneumoniae* strains and primers used in this study

| **Strains used in this study** | | | |
|---|---|---|---|
| **Strain** | **Genotype (description)**[a] | **Antibiotic resistance**[b] | **Reference or source** |
| K272 | D39 Δ*cps* Δ*pnp*::P$_c$-[*kan*$^R$-*rpsL*$^+$] (IU1945 transformed with Δ*pnp*::P$_c$-[*kan*$^R$-*rpsL*$^+$] amplicon) | Kan$^R$ | This study |
| IU1690 | D39 | None | NCTC 7466; Lanie et al., 2007 (1) |
| IU1781 | D39 *rpsL1* | Str$^R$ | Ramos-Montanez et al., 2008 (2) |
| IU1824 | D39 *rpsL1* Δ*cps2A'-cps2H'* = D39 *rpsL1* Δ*cps* | Str$^R$ | Lanie et al., 2007 (1) |
| IU1912 | D39 *spd_1717*::Tn*4001 luxABCDE* | Kan$^R$ | Ramos-Montanez et al., 2008 (2) |
| IU1918 | D39 *rpsL1 spd_1717*::Tn*4001 luxABCDE* (derived from IU1912 in Lanie et al., 2007 (1)) | Str$^R$ Kan$^R$ | Winkler lab strain |
| IU1945 | D39 Δ*cps* | None | Lanie et al., 2007 (1) |
| IU3116 | D39 *rpsL1* CEP::P$_c$-[*kan*$^R$-*rpsL*$^+$] | Kan$^R$ | Ramos-Montanez et al., 2010 (3) |
| IU4501 | D39 *rpsL1* Δ*rny*::P$_c$-[*kan*$^R$-*rpsL*$^+$] (IU1781 transformed with Δ*rny*::P$_c$-[*kan*$^R$-*rpsL*$^+$] amplicon) | Kan$^R$ | This study |
| IU4599 | D39 *rpsL1* Δ*rny* (IU4501 transformed with Δ*rny* amplicon) | Str$^R$ | This study |
| IU4867 | D39 *rpsL1* Δ*pnp*::P$_c$-[*kan*$^R$-*rpsL*$^+$] (IU1781 transformed with Δ*pnp*::P$_c$-[*kan*$^R$-*rpsL*$^+$] amplicon from K272) | Kan$^R$ | This study |
| IU4883 | D39 *rpsL1* Δ*pnp* (IU4867 transformed with Δ*pnp* amplicon) | Str$^R$ | This study |
| IU5498 | D39 *rpsL1* Δ*pnp* CEP::P$_c$-[*kan*$^R$-*rpsL*$^+$] (IU4883 transformed with CEP::P$_c$-[*kan*$^R$-*rpsL*$^+$] amplicon from IU3116) | Str$^R$ Kan$^R$ | This study |
| IU5504 | D39 *rpsL1* Δ*rny* CEP::P$_c$-[*kan*$^R$-*rpsL*$^+$] (IU4599 transformed with CEP::P$_c$-[*kan*$^R$-*rpsL*$^+$] amplicon from IU3116) | Str$^R$ Kan$^R$ | This study |
| IU6622 | D39 *rpsL1* Δ*pnp* Tn*4001 luxABCDE* (IU4883 transformed with Tn*4001 luxABCDE* amplicon from IU1912) | Str$^R$ Kan$^R$ | This study |
| IU6838 | D39 *rpsL1* Δ*rny* Tn*4001 luxABCDE* (IU4599 transformed with Tn*4001 luxABCDE* amplicon from IU1912) | Str$^R$ Kan$^R$ | This study |
| IU7058 | D39 *rpsL1* Δ*pnp*::P$_c$-[*kan*$^R$-*rpsL*$^+$] (IU4883 transformed with Δ*pnp*::P$_c$-[*kan*$^R$-*rpsL*$^+$] amplicon from K272) | Kan$^R$ | This study |

| | | | |
|---|---|---|---|
| IU7078 | D39 *rpsL1* Δ*rny*::P$_c$-[*kan*$^R$-*rpsL*$^+$] (IU4599 transformed with Δ*rny*::P$_c$-[*kan*$^R$-*rpsL*$^+$] amplicon from IU4501) | Kan$^R$ | This study |
| IU7104 | D39 *rpsL1 pnp*$^+$ (IU7058 transformed with *pnp* amplicon from IU1690) | Str$^R$ | This study |
| IU7110 | D39 *rpsL1 rny*$^+$ (IU7058 transformed with *rny* amplicon from IU1690) | Str$^R$ | This study |
| IU7152 | D39 *rpsL1 rny*$^+$ Tn*4001 luxABCDE* (IU7110 transformed with Tn*4001 luxABCDE* amplicon from IU1912) | Str$^R$ Kan$^R$ | This study |
| IU7154 | D39 *rpsL1 pnp*$^+$ Tn*4001 luxABCDE* (IU7104 transformed with Tn*4001 luxABCDE* amplicon from IU1912) | Str$^R$ Kan$^R$ | This study |
| IU10508 | D39 Δ*cps* Δ*bgaA*::*kan*-t1t2-P$_{spd_1874}$-*lacZ* | Str$^R$ Kan$^R$ | Winkler lab strain |
| NRD10025 | D39 Δ*cps rpsL1* Δ*ccnB*::P$_c$-[*kan*$^R$-*rpsL*$^+$] (IU1824 transformed with Δ*ccnB*::P$_c$-[*kan*$^R$-*rpsL*$^+$] amplicon) | Kan$^R$ | This study |
| Spn_DS_012 | D39 Δ*cps rpsL1* Δ*ccnB* (NRD10025 transformed with Δ*ccnB* amplicon) | Str$^R$ | This study |
| NRD10026 | D39 Δ*cps rpsL1* Δ*ccnD*::P$_c$-[*kan*$^R$-*rpsL*$^+$] (IU1824 transformed with Δ*ccnD*::P$_c$-[*kan*$^R$-*rpsL*$^+$] amplicon) | Kan$^R$ | This study |
| Spn_DS_013 | D39 Δ*cps rpsL1* Δ*ccnB* Δ*ccnD*::P$_c$-[*kan*$^R$-*rpsL*$^+$] (Spn_DS_012 transformed with Δ*ccnD*::P$_c$-[*kan*$^R$-*rpsL*$^+$] amplicon) | Kan$^R$ | This study |
| Spn_DS_015 | D39 Δ*cps rpsL1* Δ*ccnD* (NRD10026 transformed with Δ*ccnD* amplicon) | Str$^R$ | This study |
| NRD10027 | D39 Δ*cps rpsL1* Δ*ccnB* Δ*ccnD* (Spn_DS_013 transformed with Δ*ccnD* amplicon amplicon) | Str$^R$ | This study |
| NRD10031 | D39 Δ*cps rpsL1* Δ*ccnB* Δ*ccnD* Δ*ccnC*::P$_c$-[*kan*$^R$-*rpsL*$^+$] (NRD10027 transformed with Δ*ccnC*::P$_c$-[*kan*$^R$-*rpsL*$^+$] amplicon) | Kan$^R$ | This study |
| NRD10034 | D39 Δ*cps rpsL1* Δ*ccnB* Δ*ccnD* Δ*ccnC* (NRD10031 transformed with Δ*ccnC* amplicon) | Str$^R$ | This study |
| NRD10035 | D39 Δ*cps rpsL1* Δ*ccnB* Δ*ccnD* Δ*ccnC* Δ*ccnE*::P$_c$-[*kan*$^R$-*rpsL*$^+$] (NRD10034 transformed with Δ*ccnE*::P$_c$-[*kan*$^R$-*rpsL*$^+$] amplicon) | Kan$^R$ | This study |
| NRD10036 | D39 Δ*cps rpsL1* Δ*ccnB* Δ*ccnD* Δ*ccnC* Δ*ccnE* (NRD10035 transformed with Δ*ccnE* amplicon) | Str$^R$ | This study |
| NRD10037 | D39 Δ*cps rpsL1* Δ*ccnB* Δ*ccnD* Δ*ccnC* Δ*ccnE* Δ*ccnA*::P$_c$-[*kan*$^R$-*rpsL*$^+$] (NRD10036 transformed with Δ*ccnA*::P$_c$-[*kan*$^R$-*rpsL*$^+$] amplicon) | Kan$^R$ | This study |
| NRD10038[c] | D39 Δ*cps rpsL1 ccnB*$^+$ Δ*ccnA* Δ*ccnC* Δ*ccnD* Δ*ccnE* (NRD10037 transformed with Δ*ccnA* amplicon) | Str$^R$ | This study |
| NRD10039 | D39 Δ*cps rpsL1* Δ*bgaA*::*kan*-t1t2-P$_{VegT}$-*lacZ* | Str$^R$ Kan$^R$ | This study |
| NRD10041 | D39 Δ*cps rpsL1* Δ*bgaA*::*kan*-t1t2-P$_{VegT}$-*comC-lacZ* (IU1824 transformed with Δ*bgaA*::*kan*-t1t2-P$_{VegT}$-*comC-lacZ* amplicon) | Str$^R$ Kan$^R$ | This study |
| NRD10054 | D39 Δ*cps rpsL1 ccnB*$^+$ Δ*ccnA* Δ*ccnC* Δ*ccnD* Δ*ccnE* Δ*bgaA*::*kan*-t1t2-P$_{VegT}$-*comC-lacZ* (NRD10038 | Str$^R$ Kan$^R$ | This study |

| | | | |
|---|---|---|---|
| | transformed with Δ*bgaA*::*kan*-t1t2-$P_{VegT}$-*comC*-*lacZ* amplicon from NRD10041) | | |
| NRD10091 | D39 *rpsL1* Δ*rny*::$P_c$-[*kan*$^R$-*rpsL*$^+$] (IU1781 transformed with Δ*rny*::$P_c$-[*kan*$^R$-*rpsL*$^+$]; independently constructed using the same primers used to construct IU4501) | Kan$^R$ | This study |
| NRD10092 | D39 *rpsL1* Δ*rny* (NRD10091 transformed with Δ*rny* amplicon; independently constructed using the same primers used to construct IU4599) | Str$^R$ | This study |
| NRD10106 | D39 Δ*cps* *rpsL1* Δ*pnp*::$P_c$-[*kan*$^R$-*rpsL*$^+$] (IU1824 transformed with Δ*pnp*::$P_c$-[*kan*$^R$-*rpsL*$^+$] amplicon from K272) | Kan$^R$ | This study |
| NRD10107 | D39 Δ*cps* *rpsL1* Δ*rny*::$P_c$-[*kan*$^R$-*rpsL*$^+$] (IU1824 transformed with Δ*rny*::$P_c$-[*kan*$^R$-*rpsL*$^+$] amplicon from NRD10091) | Kan$^R$ | This study |
| NRD10108 | D39 Δ*cps* *rpsL1* Δ*pnp* (NRD10106 transformed with Δ*pnp* amplicon from IU4883) | Str$^R$ | This study |
| NRD10109 | D39 Δ*cps* *rpsL1* Δ*rny* (NRD10107 transformed with Δ*rny* amplicon from NRD10092) | Str$^R$ | This study |
| NRD10113 | D39 Δ*cps* *rpsL1* Δ*rny* Δ*bgaA*::*kan*-t1t2-$P_{VegT}$-*comC*-*lacZ* (NRD10109 transformed with Δ*bgaA*::*kan*-t1t2-$P_{VegT}$-*comC*-*lacZ* amplicon from NRD10041) | Str$^R$ Kan$^R$ | This study |
| NRD10114 | D39 Δ*cps* *rpsL1* *ccnB*$^+$ Δ*ccnA* Δ*ccnC* Δ*ccnD* Δ*ccnE* Δ*rny*::$P_c$-[*kan*$^R$-*rpsL*$^+$] (NRD10038 transformed with Δ*rny*::$P_c$-[*kan*$^R$-*rpsL*$^+$] amplicon from NRD10091) | Kan$^R$ | This study |
| NRD10117 | D39 Δ*cps* *rpsL1* *ccnB*$^+$ Δ*ccnA* Δ*ccnC* Δ*ccnD* Δ*ccnE* Δ*rny* (NRD10114 transformed with Δ*rny* amplicon from NRD10092) | Str$^R$ | This study |
| NRD10120 | D39 Δ*cps* *rpsL1* *ccnB*$^+$ Δ*ccnA* Δ*ccnC* Δ*ccnD* Δ*ccnE* Δ*rny* Δ*bgaA*::*kan*-t1t2-$P_{VegT}$-*comC*-*lacZ* (NRD10117 transformed with Δ*bgaA*::*kan*-t1t2-$P_{VegT}$-*comC*-*lacZ* amplicon from NRD10041) | Str$^R$ Kan$^R$ | This study |
| NRD10177 | D39 Δ*cps* *rpsL1* Δ*ccnD* Δ*ccnC* Δ*ccnE* Δ*ccnAB*::$P_c$-[*kan*$^R$-*rpsL*$^+$] (NRD10036 transformed with Δ*ccnAB*::$P_c$-[*kan*$^R$-*rpsL*$^+$] amplicon) | Kan$^R$ | This study |
| NRD10180 | D39 Δ*cps* *rpsL1* Δ*ccnD* Δ*ccnC* Δ*ccnE* Δ*ccnAB* (NRD10177 transformed with Δ*ccnAB* amplicon) | Str$^R$ | This study |
| NRD10187 | D39 Δ*cps* *rpsL1* Δ*ccnD* Δ*ccnC* Δ*ccnE* Δ*ccnAB* Δ*bgaA*::*kan*-t1t2-$P_{VegT}$-*comC*-*lacZ* (NRD10180 transformed with Δ*bgaA*::*kan*-t1t2-$P_{VegT}$-*comC*-*lacZ* amplicon from NRD10041) | Str$^R$ Kan$^R$ | This study |
| NRD10300 | D39 *rpsL1* Δ*pnp*::$P_c$-[*kan*$^R$-*rpsL*$^+$] (IU4883 transformed with Δ*pnp*::$P_c$-[*kan*$^R$-*rpsL*$^+$] amplicon from K272; independently constructed using the same primers used to construct IU7058) | Kan$^R$ | This study |
| NRD10302 | D39 *rpsL1* Δ*rny*::$P_c$-[*kan*$^R$-*rpsL*$^+$] (NRD10092 transformed with Δ*rny*::$P_c$-[*kan*$^R$-*rpsL*$^+$] amplicon from NRD10091; independently constructed using the same primers used to construct IU7078) | Kan$^R$ | This study |

| | | | |
|---|---|---|---|
| NRD10303 | D39 *rpsL1 pnp*$^{+}$ (NRD10300 transformed with *pnp* amplicon from IU1781; independently constructed using the same primers used to construct IU7104) | Str$^{R}$ | This study |
| NRD10305 | D39 *rpsL1 rny*$^{+}$ (NRD10302 transformed with *rny* amplicon from IU1781; independently constructed using the same primers used to construct IU7110) | Str$^{R}$ | This study |
| **Primers used for strain construction in this study** | | | |
| **Primer name** | **Primer Sequence (5'-3')** | **Template[d]** | **Product** |
| **For construction of strain K272 (Δ*pnp*::P$_{c}$-[*kan*$^{R}$-*rpsL*$^{+}$])** | | | |
| P644 | CTGAATCGAAATCAGGCTCTCCGACTCTTG | D39 | Upstream of *pnp* + 60 bp of 5’ *pnp* |
| P646 | CATTATCCATTAAAAATCAAACGGATCCTAATTATACGCCTTGCCTACAAAGATCAAGAT | | |
| Kan rpsL forward | TAGGATCCGTTTGATTTTTAATGGATAATG | P$_{c}$-[*kan-rpsL*$^{+}$] cassette[e] | P$_{c}$-[*kan-rpsL*$^{+}$] |
| Kan rpsL reverse | GGGCCCCTTTCCTTATGCTTTTG | | |
| P647 | CAAAAGCATAAGGAAAGGGGCCCCATCAAAAGGATTACAAACCTAAGAAAGAATTT | D39 | 60 bp of 3' *pnp* + downstream |
| P645 | GCATGTTAGAGAGATAGGGATCGCGATAGG | | |
| **For construction of strain IU4501, NRD10091 (Δ*rny*::P$_{c}$-[*kan*$^{R}$-*rpsL*$^{+}$])** | | | |
| DM105 | GGAAGCAGTCCTTTACTCAACATTCCGA | D39 | Upstream of *rny* + 60 bp of 5’ *rny* |
| DM103 | CACATTATCCATTAAAAATCAAACGGATCCTATCCAATGACTAAACCAATGATGACGGC | | |
| Kan rpsL forward | TAGGATCCGTTTGATTTTTAATGGATAATG | P$_{c}$-[*kan-rpsL*$^{+}$] cassette[e] | P$_{c}$-[*kan-rpsL*$^{+}$] |
| Kan rpsL reverse | GGGCCCCTTTCCTTATGCTTTTG | | |
| DM104 | CGTCCAAAAGCATAAGGAAAGGGGCCCCCAGGAAATATCAAGGTAACCGTGATTCG | D39 | 60 bp of 3' *rny* + downstream |
| DM106 | GACACGATCCACACAGAAGTGTTCTGCTTC | | |
| **For construction of strain IU4883 (Δ*pnp*)** | | | |
| P644 | CTGAATCGAAATCAGGCTCTCCGACTCTTG | D39 | Upstream of *pnp* + 60 bp of 5’ *pnp* |
| DM133 | CTTAGGTTTGTAATCCTTTTGATGACCAAGGACGTCAAAAGCAAAGT | | |
| DM134 | TACTTTGCTTTTGACGTCCTTGGTCATCAAAAGGATTACAAACCTAAGAAAGAATTTAC | D39 | 60 bp of 3' *pnp* + downstream |
| P645 | GCATGTTAGAGAGATAGGGATCGCGATAGG | | |
| **For construction of strain IU4599, NRD10092 (Δ*rny*)** | | | |
| DM105 | GGAAGCAGTCCTTTACTCAACATTCCGA | D39 | Upstream of *rny* + 60 bp of 5’ *rny* |
| DM115 | TCACGGTTACCTTGATATTTCCTGGTCCAATGACTAAACCAATGATGACGGC | | |
| DM116 | CGTCATCATTGGTTTAGTCATTGGACCAGGAAATATCAAGGTAACCGTGATTC | D39 | 60 bp of 3' *rny* + downstream |
| DM106 | GACACGATCCACACAGAAGTGTTCTGCTTC | | |
| **For construction of strain IU7104, NRD10303 (Δ*pnp* -*pnp*$^{+}$; repair strain)** | | | |
| P644 | CTGAATCGAAATCAGGCTCTCCGACTCTTG | D39 | *pnp* + flanking regions |
| P645 | GCATGTTAGAGAGATAGGGATCGCGATAGG | | |

| | | | |
|---|---|---|---|
| **For construction of strain IU7110, NRD10305 (Δ*rny* -*rny*$^{+}$; repair strain)** | | | |
| DM105 | GGAAGCAGTCCTTTACTCAACATTCCGA | D39 | *rny* + flanking regions |
| DM106 | GACACGATCCACACAGAAGTGTTCTGCTTC | | |
| **For construction of strain NRD10037 (Δ*ccnA*::P$_c$-[*kan*$^R$-*rpsL*$^{+}$])** | | | |
| Spn001 | TCATAGACAAGGCGACTGGTAAGG | D39 | Upstream of *ccnA* |
| Spn002 | CCATTAAAAATCAAACGGATCCTAAAAAAAGTT TAGGATTTTATTAAATAAAGTTAGG | | |
| Kan rpsL forward | TAGGATCCGTTTGATTTTTAATGGATAATG | P$_c$-[*kan*-*rpsL*$^{+}$] cassette[e] | P$_c$-[*kan*-*rpsL*$^{+}$] |
| Kan rpsL reverse | GGGCCCCTTTCCTTATGCTTTTG | | |
| Spn003 | TCCAAAAGCATAAGGAAAGGGGCCCGGCTTTT TGCGTGGTGAGGTGCTGGTG | D39 | Downstream of *ccnA* |
| Spn004 | GGCCCAAATAAGAGCACATGATCC | | |
| **For construction of strain NRD10025 (Δ*ccnB*::P$_c$-[*kan*$^R$-*rpsL*$^{+}$])** | | | |
| Spn005 | ATCATAGACAAGGCGACTGGTAAGGC | D39 | Upstream of *ccnB* |
| Spn006 | CCATTAAAAATCAAACGGATCCTAGGAGGTCTT TATTTAATAACTACATG | | |
| Kan rpsL forward | TAGGATCCGTTTGATTTTTAATGGATAATG | P$_c$-[*kan*-*rpsL*$^{+}$] cassette[e] | P$_c$-[*kan*-*rpsL*$^{+}$] |
| Kan rpsL reverse | GGGCCCCTTTCCTTATGCTTTTG | | |
| Spn007 | TCCAAAAGCATAAGGAAAGGGGCCCCAGGTGG AGTTTTTTAGCTCTATTTCAG | D39 | Downstream of *ccnB* |
| Spn008 | CCAAATGAACACTACGACTACCCTCACC | | |
| **For construction of strain NRD10026 (Δ*ccnD*::P$_c$-[*kan*$^R$-*rpsL*$^{+}$])** | | | |
| Spn020 | AATGAGTTAGAGCCTGGGGATGTCC | D39 | Upstream of *ccnD* |
| Spn018 | ATCCATTAAAAATCAAACGGATCCTAGAACTTA GTGTACACTCCCTAGCTTAAAGTTTCC | | |
| Kan rpsL forward | TAGGATCCGTTTGATTTTTAATGGATAATG | P$_c$-[*kan*-*rpsL*$^{+}$] cassette[e] | P$_c$-[*kan*-*rpsL*$^{+}$] |
| Kan rpsL reverse | GGGCCCCTTTCCTTATGCTTTTG | | |
| Spn019 | CGTCCAAAAGCATAAGGAAAGGGGCCCGAAAA ATGGGCTTGGTGCCTGAGAAT | D39 | Downstream of *ccnD* |
| Spn021 | GTCAGGTAATTCTCCAAGGGAATGG | | |
| **For construction of strain NRD10035 (Δ*ccnE*::P$_c$-[*kan*$^R$-*rpsL*$^{+}$])** | | | |
| Spn023 | GTATCGGTGACACCTATTTCTCTGA | D39 | Upstream of *ccnE* |
| Spn022 | CACATTATCCATTAAAAATCAAACGGATCCTAG ATTATAGTATACACATCTAATCTT | | |
| Kan rpsL forward | TAGGATCCGTTTGATTTTTAATGGATAATG | P$_c$-[*kan*-*rpsL*$^{+}$] cassette[e] | P$_c$-[*kan*-*rpsL*$^{+}$] |
| Kan rpsL reverse | GGGCCCCTTTCCTTATGCTTTTG | | |
| Spn024 | CGTCCAAAAGCATAAGGAAAGGGGCCCTCATT CATGATATAATAGAAGCAAACGGAG | D39 | Downstream of *ccnE* |
| Spn025 | CACTGATTTCAGAACTCTCTACTTCTATCC | | |
| **For construction of strain NRD10031 (Δ*ccnC*::P$_c$-[*kan*$^R$-*rpsL*$^{+}$])** | | | |
| Spn036 | GCCTTATCATATCGAGTTGGATCGCTTGC | D39 | |

| | | | |
|---|---|---|---|
| | | | Upstream of *ccnC* |
| Spn034 | CACATTATCCATTAAAAATCAAACGGATCCTAG TCTATAGTATACCCGACCTATCTTAAAC | | |
| Kan rpsL forward | TAGGATCCGTTTGATTTTTAATGGATAATG | $P_c$-[*kan-rpsL*$^+$] cassette[e] | $P_c$-[*kan-rpsL*$^+$] |
| Kan rpsL reverse | GGGCCCCTTTCCTTATGCTTTTG | | |
| Spn035 | CGTCCAAAAGCATAAGGAAAGGGGCCCGTGTT GGGATTCATGATATAATAATAAAATCG | D39 | Downstream of *ccnC* |
| Spn037 | TAATCCCCATCAATGACCCCAACTGAGT | | |
| **For construction of strain NRD10038 (Δ*ccnA*)** | | | |
| Spn013 | CAGTGGAAAAGCATGCGGAGGATTTG | D39 | Upstream of *ccnA* |
| Spn015 | TCTATCACCAGCACCTCACCACGCGACTATATA ATACTAGACCATCCT | | |
| Spn014 | AGGATGGTCTAGTATTATATAGTCGCGTGGTGA GGTGCTGGTGATAGA | D39 | Downstream of *ccnA* |
| Spn004 | GGCCCAAATAAGAGCACATGATCC | | |
| **For construction of strain Spn_DS_012 (Δ*ccnB*)** | | | |
| Spn013 | CAGTGGAAAAGCATGCGGAGGATTTG | D39 | Upstream of *ccnB* |
| Spn028 | GTCCCCAAAAGCCTGAAATAGAGCTAACTACAT GATACAAGACGAAACTTAAAAC | | |
| Spn029 | AAGTTTCGTCTTGTATCATGTAGTTAGCTCTATT TCAGGCTTTTGGGGACTATTC | D39 | Downstream of *ccnB* |
| Spn004 | GGCCCAAATAAGAGCACATGATCC | | |
| **For construction of strain NRD10034 (Δ*ccnC*)** | | | |
| Spn036 | GCCTTATCATATCGAGTTGGATCGCTTGC | D39 | Upstream of *ccnC* |
| 5'ccnCclean KO rev | CGATTTTATTATTATATCATGAATCCCAACACGT CTATAGTATACCCGACCTATCTTAAAC | | |
| 3'ccnCclean KO for | GTTTAAGATAGGTCGGGTATACTATAGACGTGT TGGGATTCATGATATAATAATAAAATCG | D39 | Downstream of *ccnC* |
| Spn037 | TAATCCCCATCAATGACCCCAACTGAGT | | |
| **For construction of strain Spn_DS_015 (Δ*ccnD*)** | | | |
| Spn020 | AATGAGTTAGAGCCTGGGGATGTCC | D39 | Upstream of *ccnD* |
| Spn031 | TTCTCAGGCACCAAGCCCATTTTTCGAACTTAG TGTACACTCCCTAGCTT | | |
| Spn030 | CTTTAAGCTAGGGAGTGTACACTAAGTTCGAAA AATGGGCTTGGTGCCTGAGAAT | D39 | Downstream of *ccnD* |
| Spn021 | GTCAGGTAATTCTCCAAGGGAATGG | | |
| **For construction of strain NRD10036 (Δ*ccnE*)** | | | |
| Spn023 | GTATCGGTGACACCTATTTCTCTGA | D39 | Upstream of *ccnE* |
| Spn033 | CCGTCCTCCGTTTGCTTCTATTATATCATGAAT GAGATTATAGTATACACATCTAATCTT | | |
| Spn032 | AAGATTAGATGTGTATACTATAATCTCATTCATG ATATAATAGAAGCAAACGGAGGACGG | D39 | Downstream of *ccnE* |
| Spn025 | CACTGATTTCAGAACTCTCTACTTCTATCC | | |
| **For construction of strain NRD10177 (Δ*ccnAB*::$P_c$-[*kan*$^R$-*rpsL*$^+$])** | | | |
| Spn001 | TCATAGACAAGGCGACTGGTAAGG | D39 | Upstream of *ccnA* |
| Spn002 | CCATTAAAAATCAAACGGATCCTAAAAAAAGTT TAGGATTTTATTAAATAAAGTTAGG | | |

| | | | |
|---|---|---|---|
| Kan rpsL forward | TAGGATCCGTTTGATTTTTAATGGATAATG | $P_c$-[*kan-rpsL*$^+$] cassette[e] | $P_c$-[*kan-rpsL*$^+$] |
| Kan rpsL reverse | GGGCCCCTTTCCTTATGCTTTTG | | |
| Spn007 | TCCAAAAGCATAAGGAAAGGGGCCCCAGGTGGAGTTTTTTAGCTCTATTTCAG | D39 | Downstream of *ccnB* |
| Spn008 | CCAAATGAACACTACGACTACCCTCACC | | |
| **For construction of strain NRD10180 (Δ*ccnAB*)** | | | |
| Spn001 | TCATAGACAAGGCGACTGGTAAGG | D39 | Upstream of *ccnA* |
| 53-ccnAB Rev2 | CCCAAAAGCCTGAAATAGAGCTGACTATATAATACTAGACCATCCTTAAATTTTGC | | |
| 53-ccnAB For2 | GCAAAATTTAAGGATGGTCTAGTATTATATAGTCAGCTCTATTTCAGGCTTTTGGG | D39 | Downstream of *ccnB* |
| Spn008 | CCAAATGAACACTACGACTACCCTCACC | | |
| **For construction of strain NRD10039 (Δ*bgaA*::*kan*-t1t2-$P_{VegT}$-*lacZ*)** | | | |
| 5'BgaFor | GATATCTGGCACTTGTCTATCACAGGTC | IU10508 | 5'*bgaA* fragment containing PvegT sequence |
| 5'Bga-T1T2-vegTrev | CAAATTATATCAAGTTAATAAGACGTTGTCAATAAAATTATTTTGACAGTAGAAACGCAAAAAGGCCATCC | | |
| 3'Bga-VegT For | ATTTTATTGACAACGTCTTATTAACTTGATATAATTTGTTTATGAAACATCTTGATCCCGTCGTTTTAC | IU10508 | 3'*bgaA* fragment containing PvegT sequence |
| 3'Bga-rpsLKan Rev | CTGGTTTTTCCTTAGTCAACTGGATACGG | | |
| **For construction of strain NRD10041 (Δ*bgaA*::*kan*-t1t2-$P_{VegT}$-*comC-lacZ*)** | | | |
| 5'BgaFor | GATATCTGGCACTTGTCTATCACAGGTC | NRD10039 | 5'*bgaA* fragment |
| 5'VegT-comC-Rev | ATGTTTTTTCTTGTAAGCTAACTTACAAACAAATTATATCAAGTTAATAAGACGTTGTCA | | |
| mVegT-comC-For | TCTTATTAACTTGATATAATTTGTTTGTAAGTTAGCTTACAAGAAAAAACATTTTAGGAG | D39 | PvegT + *comC* 5'UTR and first 11 codons |
| mVegT-comC-Rev | CCCAGTCACGACGTTGTAAAACGACAGCTACAAACTGTTCCAATTTAACTGTGTTTTTC | | |
| 3'VegT-comC-For | GAAAAACACAGTTAAATTGGAACAGTTTGTAGCTGTCGTTTTACAACGTCGTGACTGGG | NRD10039 | 3'*bgaA* fragment |
| 3'Bga-rpsLKan Rev | CTGGTTTTTCCTTAGTCAACTGGATACGG | | |

[a]Strains were constructed by transformation of fusion PCR amplicons into the indicated recipient strain as described in Materials and Methods.

[b]Antibiotic resistance markers: Kan, kanamycin; Str, streptomycin.

[c]Transformation of the Δ*ccnA* amplicon into NRD10037 background made the resulting strain *ccnB*$^+$.

[d]Genomic DNA of indicated *S. pneumoniae* strains was used as templates for PCR reactions, except for $P_c$-[*kan-rpsL*$^+$].

[e]$P_c$-[*kan-rpsL*$^+$] cassettes are described in (4).

[f]$P_{vegT}$ sequence is derived from *B. subtilis* as described in (5).

**Table S2.** Growth characteristics of Δ*rny* mutants used in this study

| Strain | Genotype | Doubling time (min)[a,c] | Growth yield ($OD_{620}$)[b,c] | Reference |
|---|---|---|---|---|
| **Growth in 5-day old BHI at 37$^O$C** | | | | |
| IU1781 | WT (D39 *rpsL1*) | 37.5 ± 0.3 | 0.95 ± 0.01 | Fig. S1A |
| NRD10092 | Δ*rny* | 55.0 ± 6.2* | 0.32 ± 0.04**** | |
| NRD10305 | Δ*rny*//*rny*$^+$ | 40.1 ± 0.8 $^{ns}$ | 0.94 ± 0.06 $^{ns}$ | |
| **Growth in 15-day old BHI at 37$^O$C** | | | | |
| IU1781 | WT (D39 *rpsL1*) | 44.6 ± 2.0 | 0.83 ± 0.01 | Fig. 1A |
| NRD10092 | Δ*rny* | 76.0 ± 9.6* | 0.13 ± 0.01**** | |
| NRD10305 | Δ*rny*//*rny*$^+$ | 51.2 ± 1.0$^{ns}$ | 0.78 ± 0.01* | |
| **Growth in 25-day old BHI at 37$^O$C** | | | | |
| IU1781 | WT (D39 *rpsL1*) | 46.4 ± 0.8 | 0.75 ± 0.02 | Fig. S1B |
| NRD10092 | Δ*rny* | 65.9 ± 4.8** | 0.11 ± 0.01**** | |
| NRD10305 | Δ*rny*//*rny*$^+$ | 48.9 ± 0.7 $^{ns}$ | 0.72 ± 0.03 $^{ns}$ | |
| **Growth in 3-day old BHI at 32$^O$C** | | | | |
| IU1781 | WT (D39 *rpsL1*) | 54.4 ± 0.2 | 0.95 ± 0.003 | Fig. 1B |
| NRD10092 | Δ*rny* | 77.0 ± 0.2**** | 0.12 ± 0.01**** | |
| NRD10305 | Δ*rny*//*rny*$^+$ | 57.6 ± 0.4$^{ns}$ | 0.86 ± 0.01*** | |
| **Growth in 15-day old BHI at 37$^O$C** | | | | |
| IU1781 | WT (D39 *rpsL1*) | 63.5 ± 0.7 | 0.63 ± 0.01 | Fig. S1C |
| NRD10092 | Δ*rny* | 88.3 ± 4.8** | 0.09 ± 0.01**** | |
| NRD10305 | Δ*rny*//*rny*$^+$ | 67.3 ± 0.8 $^{ns}$ | 0.38 ± 0.02**** | |

[a]Doubling times were determined from data points in the exponential growth phase indicated by semi-log plots of $OD_{620}$ vs time.

[b]Maximum growth yield is the highest $OD_{620}$ value obtained during a 7 to 10 h period after subculture to a starting inoculum of $OD_{620}$ ≈ 0.002 in BHI broth.

[c]Mean values of doubling times and growth yields are presented with ± SEM and were calculated using GraphPad Prism version 7. ns, *,**,***,**** indicate that the values are nonsignificant, or P values are < 0.05, 0.01, 0.001, or 0.0001 respectively, when compared with the strain in the first line of each section using ANOVA with GraphPad Prism 7.

**Table S3.** Droplet-Digital PCR primers and oligonucleotide probes used in this study

| **Droplet-Digital PCR primers used in this study** | | |
|---|---|---|
| **Primer name** | **Primer Sequence (5'-3')** | **Gene** |
| qmapZ-for | GTGAACATACGCTTGCCAAATC | *mapZ* |
| qmapZ-rev | TCCAACACACCATCCACAATAG | |
| qclpL-for | TGTGGAACGAATGACAGGTATC | *clpL* |
| qclpL-rev | TACGGCCTTATCTTGACCAATC | |
| qdnaK-for | ACCCTGATGAAGTAGTTGCTATG | *dnaK* |
| qdnaK-rev | CAAGTGACAATGGCGTTACATC | |
| qspd_0703-for | GAATTAAACCAGCAGCTTCCTC | *spd_0703* |
| qspd_0703-rev | GATTTCCTTGAGATTGGCATCAG | |
| qspd_1604-for | GAAGAGTATAAAGGTGTACCGAGATG | *spd_1604* |
| qspd_1604-rev | GCTTCTAGGTAGTCTGCAATCG | |
| qtrpE-for | CCGCATATCGCTCCACATAA | *trpE* |
| qtrpE-rev | GAAACGCTGGCTGAGTACA | |
| KK461 | CTGGCTTGAACGGAACAACCAAGA | *trpD* |
| KK462 | CCAGCATTCAAGACTGTCGTTTCC | |
| qtrpA-for | GCGCCAGAAAGAGTTGATTG | *trpA* |
| qtrpA-rev | GTCCAAATCTGCACGGTAATTG | |
| qalaS-for | CAGAAGGAACTGCCTCTCTTATC | *alaS* |
| qalaS-rev | TCACAGTAGCCACAATCTTACC | |
| qspd_0437-for | GCCATCTCATTTGTGCTGATG | *spd_0437* |
| qspd_0437-rev | CACAAGGACTGGCAAGATACTA | |
| KK489 | CAGCAGTAGGGAATCTTCGGCAAT | *16S rRNA* |
| KK490 | TACGCCCAATAAATCCGGACAACG | |

| **Oligonucleotide probes used in this study** | |
|---|---|
| **sRNA ID** | **Oligonucleotide probe sequence in 5′-3′ direction** |
| CcnA | ACCTGATTGGGTGGCTTCATTAGGAGATTGTGATG |
| CcnB | CTCCACCTGATTGGGTGGAGTTAAGGGAGATTATTATG |
| CcnC | TTTGTTACAACAAGTTAGGAGGTCTTCTTGTAACC |
| CcnD | GGGAGATTATTATGAAAAAGTTTTAGGAGTTTAAGTTAAGG |
| CcnE | TAAAAGCCACCCATACAGGCGACTTTTGAAGGAG |
| Spd_sr12-1 | GGAATTTCCCTATCGCTCAGTCTCGCAATAACGAG |
| Spd_sr12-2 | CATTGCCACTTATCTGTTTTCAGCTACCACAGACTCTC |
| Spd_sr20 | GCCCCATATGACCTATAATGAAAAGCGTCTAACCAACTC |
| Spd_sr32 | AACCAGTTTTGACACATTTGTGGACTTCTCAGCG |
| Spd_sr43 | CCCTTCGCCAGTCTTAACTGTATCAGGTTCAATGGG |

| | |
|---|---|
| Spd_sr54 | GATACGTCTTGTCCTCGGCAGGATATTTATGAGTCG |
| Spd_sr70 | CTCTGGACTAAGACAAGTGAAAATCAATTCTCAAC |
| Spd_sr73 | AAGAGTAAACTCAGCTAGTCCAACTAACTGAGTT |
| Spd_sr74 | CTACAGAAAGCGCCAGCCCTTTATTTTGCCCTACT |
| Spd_sr80 | GAGGGACACCTCTGATCGGCTCTAACGTGGCCACC |
| Spd_sr82 | CATTATCACACCTTTCTAAGGTGGTTTTTTTATCCCGT |
| Spd_sr83 | TTACTTTAATCGTTACTGTCATATGAGAGTCCTCG |
| Spd_sr88 | CCAACCACCTATTCACAATCACCACAGGCTCCC |
| Spd_sr96 | GATTGCTCCAGTAATAAAACCATTGGGAATGAAGG |
| Spd_sr100 | TTCTCTGTGTGTAGTGTACTTGCCACAATGCTTAC |
| Spd_sr108 | GAAAATCAAAGTGCAAACTAGGAAGCTAGCCGCAG |
| Spd_sr114 | GTGCGGTCGAAGTTTAACACTTCCTCTCAGACTGT |
| Spd_sr116 | ACTTCAACTCGCAAAACGACCCAAGGGCAACTGCTT |

**Table S4:** Half-life measurements

| sRNA | Strain[a] | Half-life[b] (min) |
|---|---|---|
| CcnA | WT | 17.6 ± 1.7 |
| | Δ*rny* | 52.2 ± 15 |
| CcnB | WT | 14.1 ± 1.7 |
| | Δ*rny* | 59.9 ± 15 |
| CcnC | WT | 6.0 ± 1.4 |
| | Δ*rny* | 22.9 ± 4.1 |
| CcnE | WT | 15.8 ± 1.5 |
| | Δ*rny* | 28.4 ± 3.4 |

[a]RNA stability for CcnA, CcnB, CcnC and CcnE in the D39 parent strain (IU1781) and isogenic mutant Δ*rny* mutant (NRD10092) were determined as described in Materials and Methods.

[b]Each half-life measurement represents the average of at least three independent determinations. Corresponding data are graphed in Figure 5 and S6.

### SUPPLEMENTAL FIGURES

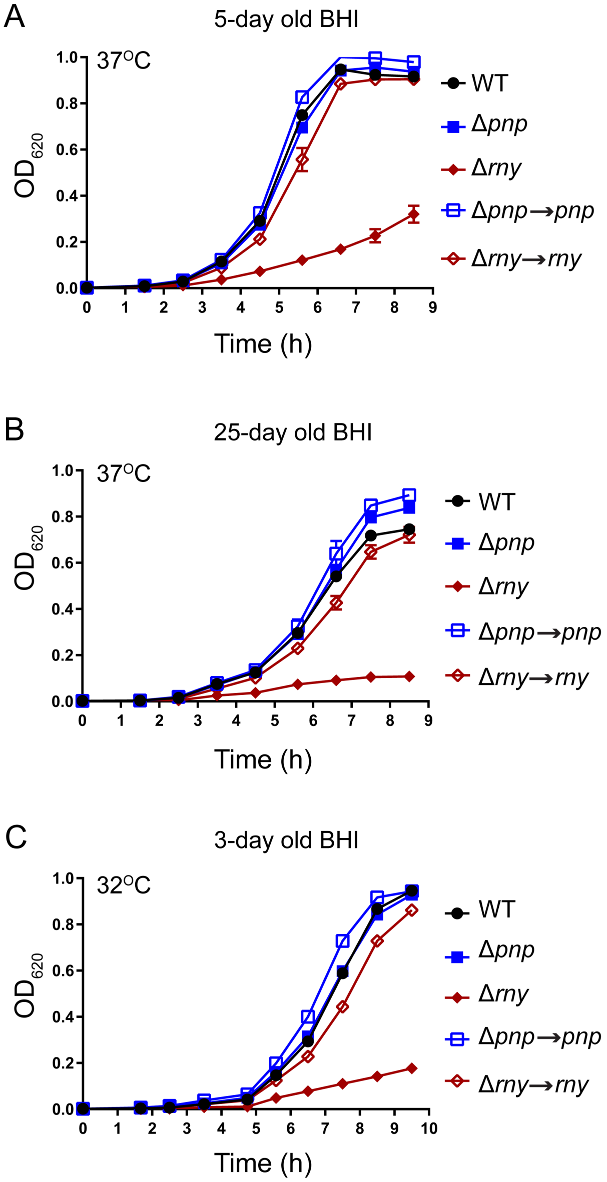

**Figure S1. Growth curves of Δ*rny* and Δ*pnp* mutants in batches of BHI broth of different ages at optimal (37°C) and lower (32°C) temperatures.** Growth curves of encapsulated D39 parent strain (IU1781) and isogenic mutant strains, Δ*rny* (NRD10092), Δ*pnp* (IU4883), Δ*rny//rny*$^{+}$ (NRD10305), and Δ*pnp//pnp*$^{+}$ (NRD10303), in 5-day-old- (A) or 25-day-old (B) BHI broth at 37°C or 3-day-old BHI broth at 32°C (C). Points and error bars (where not visible, error bars are smaller than the symbol) represent the means and standard errors (SEM) of the growth curves for three independent replicates for each strain tested. Average growth rates and growth yields are listed in Table S2.

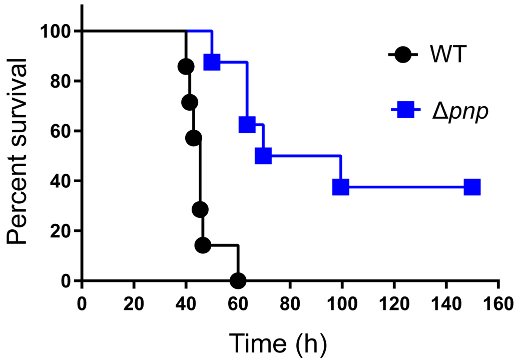

**Figure S2. Survival curve analysis showing disease progression of Δ*pnp* mutant compared to a WT parent in a murine pneumonia model.** Eight ICR male mice were inoculated intranasally with ≈$10^7$ CFUs in 50 µL inocula of the D39 parent (IU1781) strain or an isogenic Δ*pnp* mutant (IU4883), and disease progression was followed in real time by a survival curve analysis (*Materials and Methods*). Survival curves were analyzed by Kaplan-Meier statistics and log-rank tests to determine P-values.

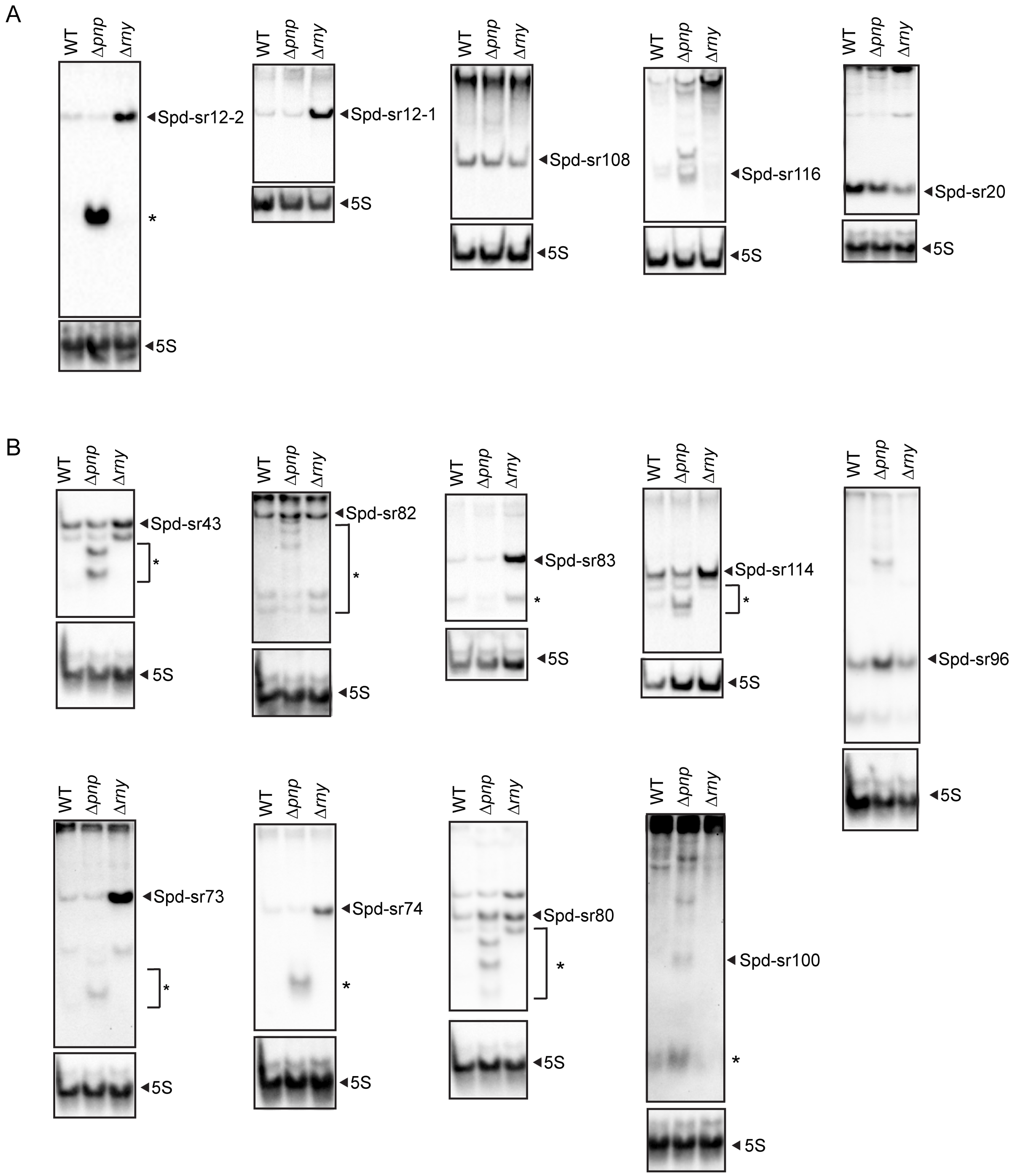

**Figure S3. Northern blot validations for sRNA-seq analysis.** (A) Northern blot validations for sRNAs that showed differential expression in a Δ*rny* mutant (NRD10092) relative to the WT parent (IU1781) parent in sRNA-seq analysis (see Table 3). Changes in Spd-sr20 amount were not detected by sRNA-seq (Table 3). The levels of the sRNAs were also determined in an isogenic Δ*pnp* mutant (IU4883). (B) Northern blot validations for sRNAs that showed differential expression in a Δ*pnp* mutant relative to the WT parent (IU1781) in sRNA-seq analysis (see Table 4). Changes in Spd-sr96 amount were not detected by RNA-seq analysis (Table 4), but its levels were still examined by northern blotting. The levels of the sRNAs were also determined in an isogenic Δ*rny* mutant. Black triangles and asterisk (*) indicate the full-length sRNA transcripts and sRNA decay products, respectively.

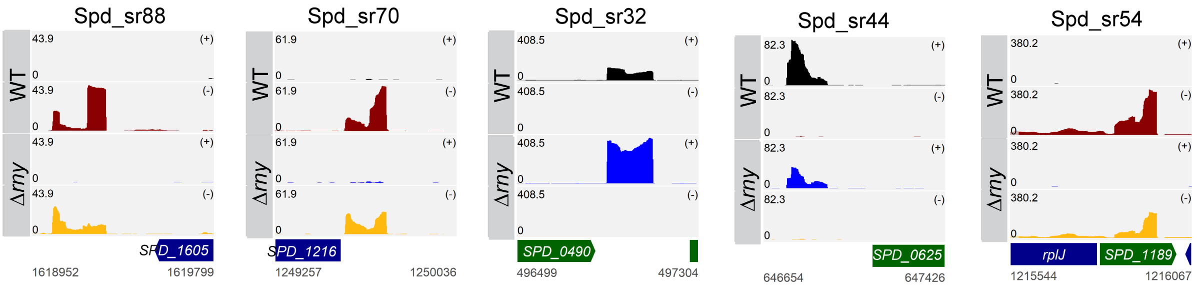

**Figure S4. Read coverage maps for sRNAs shown in Figure 4 in the Δ*rny* mutant compared to the WT parent.** Tracks labeled WT and Δ*rny* correspond to the read coverages for the sRNAs and their flanking regions in the WT (IU1781) and Δ*rny* (NRD10092) mutant strain, respectively, obtained from sRNA-seq. Coverage represents depth per million reads of paired-end sRNA fragments from averaged normalized replicates (see Materials and Methods). In each coverage graph, ORFs encoded on the plus and the minus strands are color-coded in green and blue, respectively.

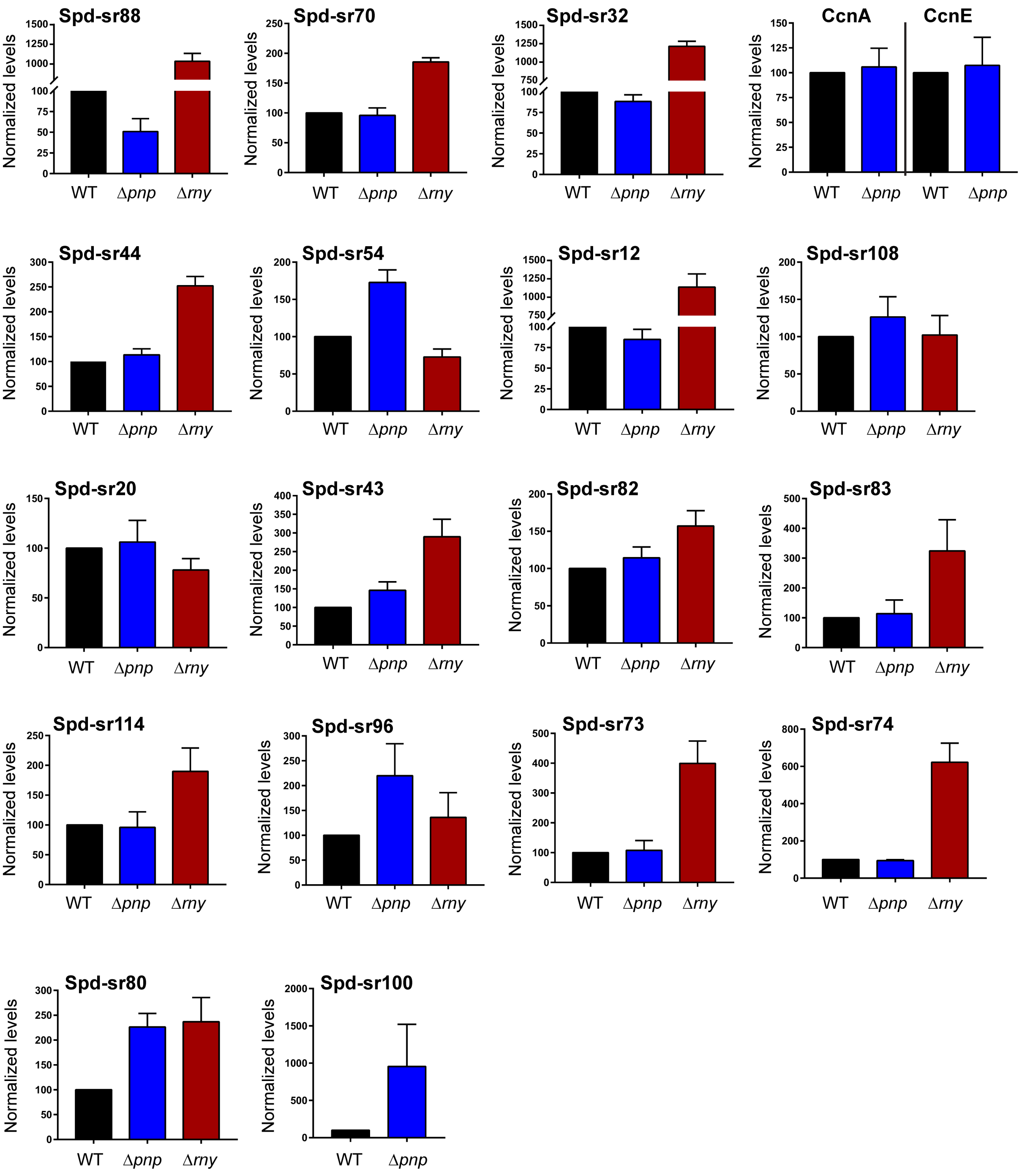

**Figure S5. Corresponding northern blot quantifications for Figure 4 and Figure S3.** sRNA steady-state levels were determined from northern blots of RNA isolated from exponentially growing cultures of the Δ*pnp* mutant (IU4883), Δ*rny* mutant (NRD10092) or WT parent (IU1781)

strain as described in the Materials and Methods. Signal intensities of the full-length sRNA transcripts were quantified using northern blotting and normalized to their corresponding loading controls (5S rRNA). The normalized value for each sRNA isolated from the WT strain was set at 100, and levels of sRNA in each mutant are scaled to this value. The full-length transcript for Spd-sr100 was only detected in a Δ*pnp* mutant (see Figure S3).

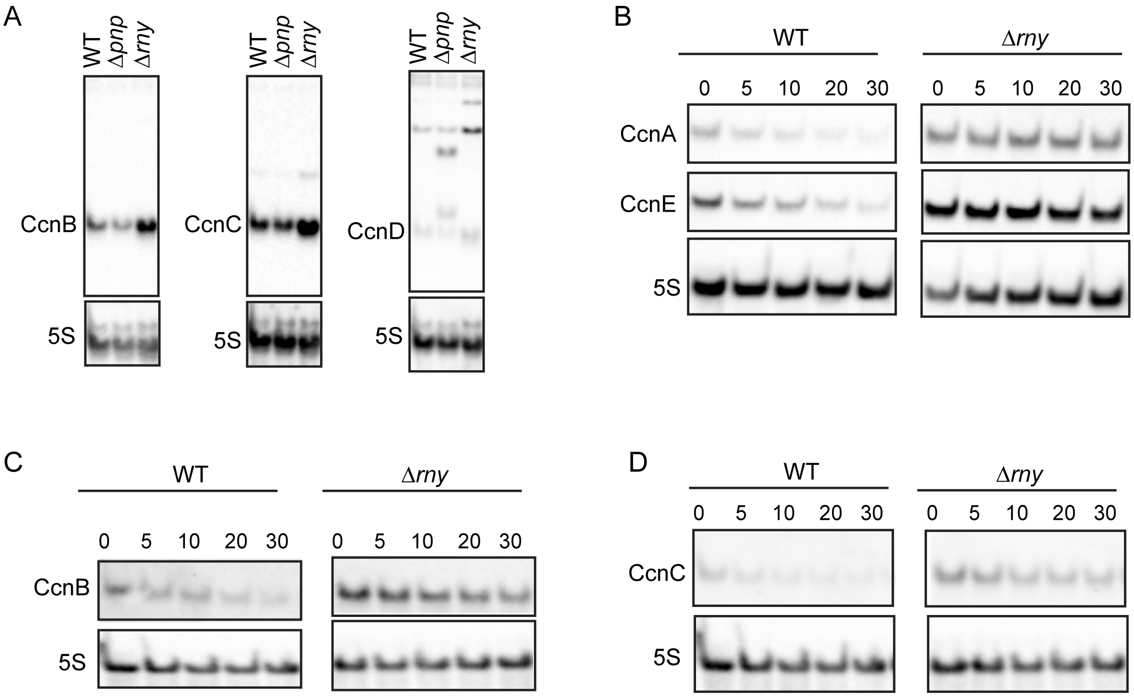

**Figure S6.** Northern blots corresponding to data presented in Figure 5. (A) Representative northern blots corresponding to CcnB, CcnC, and CcnD steady-state levels in the Δ*rny* mutant compared to a WT strain shown in Figure 5G. (B-D) Representative northern blots corresponding to CcnA, CcnB, CcnC, and CcnE stability curves shown in Figure 5C, 5F, 5H and 5I, respectively. Northern blot analysis was used to determine CcnA, CcnB, CcnC and CcnE and 5S rRNA (loading control) amounts at the indicated time points following rifampicin addition to the WT and the Δ*rny* mutant that were growing exponentially as described in Material and Methods.
